# A kinesin-1 variant reveals motor-induced microtubule damage in cells

**DOI:** 10.1101/2021.10.19.464974

**Authors:** Breane G. Budaitis, Somayesadat Badieyan, Yang Yue, T. Lynne Blasius, Dana N. Reinemann, Matthew J. Lang, Michael A. Cianfrocco, Kristen J. Verhey

## Abstract

Kinesins drive the transport of cellular cargoes as they walk along microtubule tracks, however, recent work has suggested that the physical act of kinesins walking along microtubules can stress the microtubule lattice. Here, we describe a kinesin-1 KIF5C mutant with an increased ability to generate defects in the microtubule lattice as compared to the wild-type motor. Expression of the mutant motor in cultured cells resulted in microtubule breakage and fragmentation, suggesting that kinesin-1 variants with increased damage activity would have been selected against during evolution. The increased ability to damage microtubules is not due to the altered motility properties of the mutant motor as expression of the kinesin-3 motor KIF1A, which has similar single-motor motility properties, also caused increased microtubule pausing, bending, and buckling but not breakage. In cells, motor-induced microtubule breakage could not be prevented by increased a-tubulin K40 acetylation, a post-translational modification known to increase microtubule flexibility. *In vitro*, lattice damage induced by wild-type KIF5C was repaired by soluble tubulin and resulted in increased rescues and microtubule growth whereas lattice damage induced by the KIF5C mutant resulted in larger repair sites that made the microtubule vulnerable to breakage and fragmentation when under mechanical stress. These results demonstrate that kinesin-1 motility causes defects in and damage to the microtubule lattice in cells. While cells have the capacity to repair lattice damage, conditions that exceed this capacity result in microtubule breakage and fragmentation and may contribute to human disease.

## Introduction

Kinesins are a superfamily of proteins that carry out diverse microtubule-dependent processes such as cell division, cell motility, axonal transport, and cilium assembly and function (Hirokawa et al., 2009; 2015). Kinesin proteins are defined by the presence of a kinesin motor domain which contains sequences for nucleotide and microtubule binding. The conventional kinesin function of cargo transport involves processive motility wherein the kinesin motor domain converts the energy of ATP hydrolysis into force and directed motion along the microtubule surface. Much work has focused on understanding the molecular mechanisms by which kinesins generate processive motility and has revealed critical contributions for motor protein dimerization, ATP binding and hydrolysis, and conformational changes of the neck linker (NL) [reviewed in [1-6]].

Recent work has raised the possibility that the physical act of motors walking on microtubules creates stress in the microtubule lattice. In microtubule gliding assays, high densities of kinesin-1 motors can cause breakage of taxol-stabilized microtubules and their splitting into protofilaments [7-11]. Cryo-electron and fluorescence microscopy analyses revealed that the kinesin-1 motor domain, when in its strong microtubule-binding state (ATP-bound or apo), induces a change in the conformation of tubulin subunits within the GDP lattice [12-15]. This conformational change can manifest to adjacent tubulin subunits and positively influence subsequent kinesin binding events in the same region of the microtubule [15, 16].

Microtubule gliding assays involve motors working in teams while anchored to a solid substrate and it has been unclear whether the processive motility of single kinesin motors walking on the microtubule could also create stress on the lattice. Recent work using *in vitro* motility assays demonstrated that motility of the mammalian kinesin-1 motor KIF5B, the yeast kinesin-8 motor Klp3, and yeast cytoplasmic dynein results in microtubule destruction via breakage and depolymerization [17]. Although it was widely known that members of the kinesin-8 and kinesin-13 families can catalyze conformational changes to tubulin subunits that result in filament disassembly at the ends of microtubules [18, 19], this was the first demonstration of motor stepping-induced destruction occurring along the shaft of the microtubule.

While these findings have implications for the stability and function of microtubule networks in cells [20], whether processive motility of kinesin and/or dynein motors creates stress and/or defects in the microtubule lattice in cells has not been determined. Here, we describe a kinesin-1 mutant that causes microtubule destruction when expressed in cells. The identification of this destructive kinesin-1 mutant arose from our recent work investigating the role of the NL in force generation of kinesin-1 and kinesin-3 motors [21-24]. The NL is a short and flexible segment that links the kinesin motor domain to the first coiled coil (the neck coil) that drives dimerization of kinesin proteins [25, 26]. We generated a series of truncations in the rat kinesin-1 motor KIF5C and then focused on a variant that lacks half of the coverstrand (deletion of amino acids 2-6, hereafter referred to as Δ6). Using *in vitro* single-molecule assays, we found that Δ6 motors display reduced force generation (stall force ∼1 pN) but enhanced motility properties (velocity, processivity, landing rate). These results support the hypothesis that the strength of NL docking involves a tradeoff between speed and force generation [22, 24, 27].

Surprisingly, we found that expression of Δ6 motors resulted in destruction of the microtubule network in cells. Using *in vitro* assays, we show that Δ6 motors generate increased microtubule destruction as compared to the wild-type kinesin-1. Although soluble tubulin can repair lattice damage induced by both wild-type and mutant motors, the increased damage caused by Δ6 results in microtubule breakage and destruction. These findings suggest that the mutant is an unnatural or rogue motor whose activity would have been selected against during evolution. These results also indicate that cells must repair lattice defects to prevent microtubule breakage and fragmentation as motor-induced damage results in microtubules that are unable to bear compressive loads.

## Results

### Truncation of the kinesin-1 coverstrand results in reduced force output but enhanced motility

During force generation, the NL responds to the nucleotide state of the motor domain to dock along the side of the motor domain in two steps. The first step, which we term zippering, occurs in response to ATP binding and involves zippering of the first half of the NL (the β9 segment) to the coverstrand (the β0 segment) to form a 2 stranded β-sheet called the cover-neck bundle (CNB). The second step, which we term latching, involves binding of the second half of the NL (the β10 segment) within a docking pocket to latch the NL in place (Figure S1). Initial work describing a role for the coverstrand in CNB formation and force generation noted that the length of the coverstrand varies across kinesin families [21]. Evidence that the length of the coverstrand is critical for kinesin-1 motility and force generation comes from two studies where (i) deletion of the entire N-terminal coverstrand of *Drosophila* kinesin-1 resulted in severely stunted motility and minimal force generation [22] and (ii) replacement with the shorter coverstrand of human kinesin-5 resulted in milder defects in velocity, processivity, and stall force [27]. To examine how the length of the kinesin-1 coverstrand influences CNB formation and force generation, we created a series of truncations that successively remove single residues from the N-terminus and thereby shorten the coverstrand (Figure S1D). All truncations were generated in the context of a constitutively-active version of rat KIF5C containing aa 1-560 [*Rn*KIF5C(1-560)]. In initial experiments, all truncations displayed similar increases in velocity and processivity in single-molecule motility assays and thus, for the purposes of brevity and clarity, only the results of truncation KIF5C(1-560)-Δ6, which removes half of the coverstrand (Figure 1A, Figure S1B,D), will be reported here.

**Figure 1.**
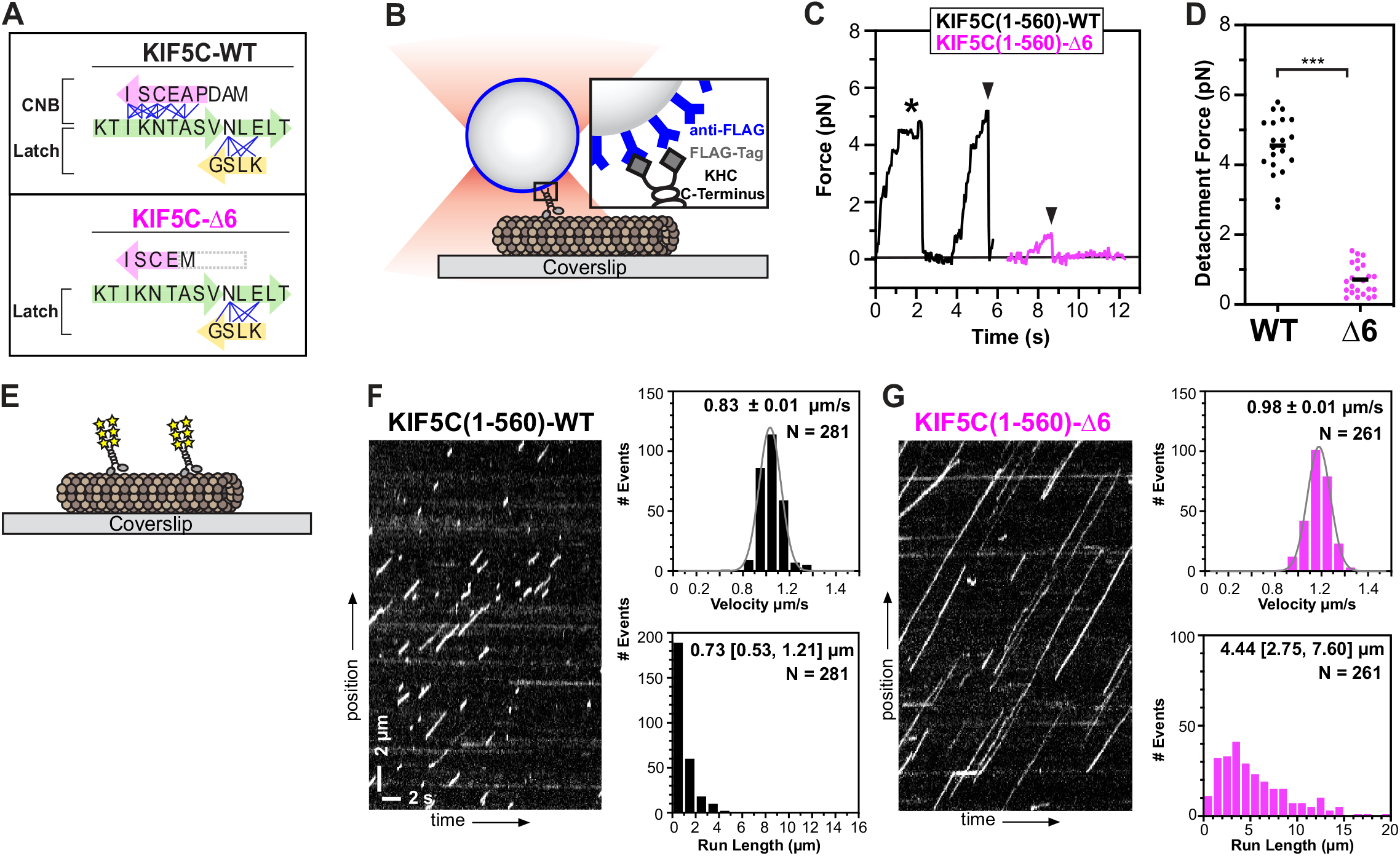
Truncation of the coverstrand results in reduced kinesin-1 force output but enhanced motility. (A) Cartoon schematic of key structural elements involved in kinesin-1 NL docking. The first half of the NL (β9, green) interacts with the coverstrand (β0, magenta) to form the cover-neck bundle (CNB). The second half of the NL (β10, green) interacts with β7 (yellow) of the core motor domain for NL latching. Residue-residue contacts for NL docking are depicted as blue lines (Budaitis et al., 2019). The coverstrand residues missing in the Δ6 truncation are indicated a grey box. (B-D) Motility under load. (B) Schematic of single-molecule optical trap assay. Flag-tagged KIF5C(1-560)-WT or KIF5C(1-560)-Δ6 motors were attached to beads functionalized with anti-Flag antibodies and subjected to standard optical trap assays. (C) Representative force versus time records of beads driven by single KIF5C(1-560)-WT (black) or KIF5C(1-560)-Δ6 (magenta) motors. The asterisk indicates a stall event; black arrowheads indicate abrupt detachment events. (D) Detachment forces. Each dot indicates the detachment force of a motility event and the black line indicates the mean value for the population. N > 20 events for each construct; ***, p<0.001 (Student’s t-test). (E-G) Motility under unloaded conditions. (E) Schematic of unloaded, single-molecule motility assay. The motility of 3xmCit-tagged KIF5C(1-560)-WT or KIF5C(1-560)-Δ6 motors along taxol-stabilized microtubules was determined in standard single-molecule motility assays using TIRF microscopy. (F,G) Representative kymographs with time displayed on the x-axis (bar 2s) and distance displayed on the y-axis (bar 2 μm). From the kymographs, single motor velocities and run lengths were determined. Population data for motor velocities and run lengths are plotted as histograms with the mean +/-SEM or median [quartiles], respectively, indicated at the top. N=281 or 261 motility events across three independent experiments.

We used a custom-built optical trap apparatus with nanometer-level spatial resolution to assess the effect of the Δ6 coverstrand truncation on kinesin-1’s force output. COS-7 cell lysates containing Flag-tagged KIF5C(1-560)-WT or KIF5C(1-560)-Δ6 kinesin-1 motors were bound to anti-Flag-coated beads and subjected to standard single-molecule trapping assays (Figure 1B) [24, 28, 29]. Individual KIF5C(1-560)-WT motors were motile in the absence of load, and upon applying the laser trap, stalled on the microtubule when approaching the detachment force, and detached from the microtubule at an average force of 4.6 ±0.8 pN (Figure 1C,D), consistent with previous studies [22, 24, 27, 30]. In contrast, KIF5C(1-560)-Δ6 mutant motors often detached from the microtubule before stalling (Figure 1C) and at much lower loads than WT motors with a mean detachment force of 0.7 ± 0.4 pN (Figure 1D). The reduced force output of the KIF5C(1-560)-Δ6 protein is similar to that observed previously for kinesin-1 motors with point mutations in the coverstrand that impair CNB formation [22, 24].

We then examined the motility properties of KIF5C(1-560)-Δ6 mutant motors under unloaded conditions. COS-7 cell lysates containing KIF5C(1-560)-WT or KIF5C(1-560)-Δ6 motors tagged with three tandem monomeric citrine fluorescent proteins (3xmCit) were added to flow chambers containing taxol-stabilized microtubules and their single-molecule motility was examined using total internal reflection fluorescence (TIRF) microscopy (Figure 1E). The velocities and run lengths were determined from kymograph analysis and summarized as a histogram. Individual KIF5C(1-560)-WT motors underwent directed motility with speed (average 0.83 ± 0.01 μm/s) and processivity (median 0.73 μm [quartiles 0.53, 1.21 μm]) (Figure 1F), comparable to previous work [22, 24]. In contrast, individual KIF5C(1-560)-Δ6 motors were faster (average 0.98 ± 0.01 μm/s) and more processive (median 4.4 μm [quartiles 2.75, 7.60 μm]) (Figure 1G). The enhanced motility properties of the KIF5C(1-560)-Δ6 motor are similar to those previous observed for kinesin-1 motors with point mutations in the coverstrand that impair CNB formation [24], suggesting that mutations which shorten and/or impair CNB formation are tolerated by the motor when stepping under no load.

Collectively, these results support the model that the coverstrand plays a critical mechanical role for single kinesin motors to step under load. These results also highlight how subtle changes in the coverstrand can act as a molecular gear shift, where motor speed and processivity come at the cost of robust force production [4, 21-24, 27, 31].

### Expression of Δ6 mutant motors results in destruction of the microtubule network in cells

We next set out to test whether the coverstrand truncation Δ6 impacts the ability of kinesin motors to work in teams to drive cargo transport in cells. For this, we aimed to employ organelle dispersion assays (Figure S2A) as utilized previously [24, 32, 33], however, we were surprised to find that COS-7 cells expressing KIF5C(1-560)-Δ6 mutant motors had dispersed Golgi elements in the absence of motor recruitment (Figure S2B). We also noticed that the KIF5C(1-560)-Δ6 mutant motors decorated highly curved and knotted microtubules in cells (Figure S2B). We thus considered the possibility that expression of KIF5C(1-560)-Δ6 mutant motors may cause unexpected changes to the underlying microtubule network in cells.

To test this hypothesis, we compared the organization of the microtubule network in COS-7 cells expressing the KIF5C(1-560)-Δ6 mutant motor to cells expressing the KIF5C(1-560)-WT motor (Figure 2A,B). All expressed motors were tagged with 3xmCit at their C-termini. We quantified the organization of the microtubule network using three different parameters. First, to quantify microtubule destruction, the number of microtubule fragments per cell was determined and plotted against motor expression level on a scatter plot (Figure 2D). Second, to quantify loss of the microtubule network at the cell periphery, the total length of the microtubule network in a 100 × 100 pixel box at the cell periphery was determined and plotted against motor expression level on a scatter plot (Figure 2E). Third, to quantify the density of the microtubule network at the cell periphery, we determined the number and size (area) of the pores between microtubules (*i*.*e*., the porosity of the network) in a 100 × 100 pixel box at the cell periphery (Figure 2F).

**Figure 2.**
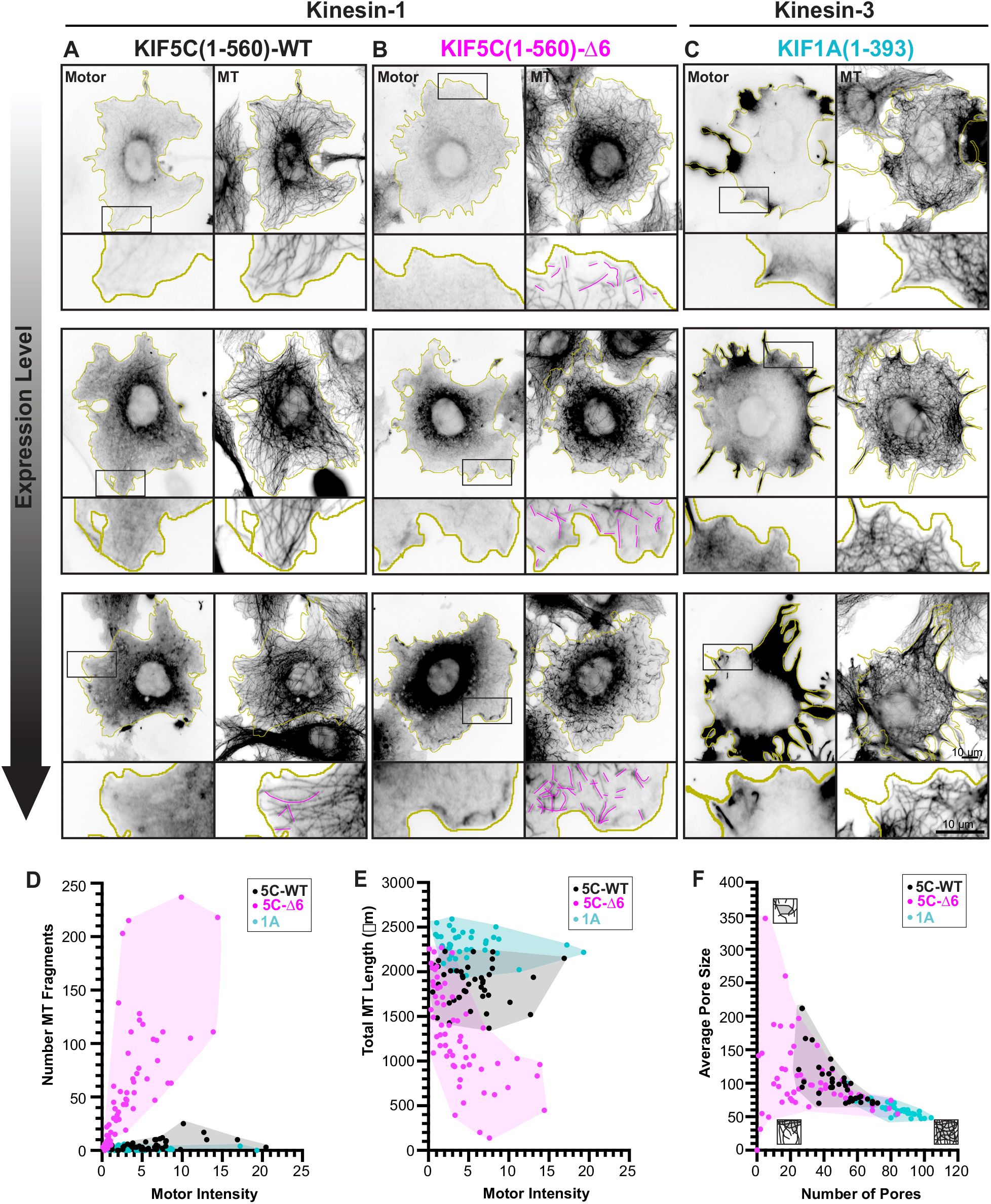
Expression of KIF5C(1-560)-Δ6 results in microtubule destruction in cells. (A-C) Representative images of the microtubule (MT) network in cells expressing low, medium, or high levels of motor. COS-7 cells were transfected with plasmids encoding for the expression of 3xmCit-tagged (A) KIF5C(1-560)-WT, (B) KIF5C(1-560)-Δ6, or (C) KIF1A(1-393) motors and then fixed and stained with an antibody against α-tubulin. Yellow lines indicate the periphery of each cell; black boxes indicate regions shown below each image at higher magnification; magenta traces indicate microtubule fragments. Scale bars, 10 μm. (D-F) Quantification of changes in the microtubule network. Each dot represents one cell [black, KIF5C(1-560)-WT; magenta, KIF5C(1-560)-Δ6; cyan, KIF1A(1-393)]. N > 39 cells each across 2 independent trials. (D) Number of microtubule fragments versus motor expression level. (E) Total length of microtubules in a 100×100 pixel box at the cell periphery versus motor expression level. (F) Density of microtubules in a 100×100 pixel box at the cell periphery as described by the number and size of the pores in the microtubule network. Cartoons depict the density of the microtubule network at each extreme (microtubules are black, pores in the network are grey).

The microtubule network in cells expressing kinesin-1 KIF5C(1-560)-WT displayed a characteristic radial array that extends to the cell periphery, with a small number of microtubule fragments observed in cells expressing very high levels of the WT motor (Figure 2A, magenta lines). In contrast, the microtubule network in cells expressing kinesin-1 KIF5C(1-560)-Δ6 mutant motors was dramatically disrupted. Microtubule fragments were observed at the periphery even in cells expressing low levels of KIF5C(1-560)-Δ6 motors (Figure 2B magenta lines, Figure 2D). Cells expressing the KIF5C(1-560)-Δ6 mutant motors also displayed a dramatic loss of microtubules at the cell periphery (Figure 2B,E) and a corresponding increase in the porosity of the microtubules in the cell periphery (Figure 2B,F).

To test whether it is the altered motility properties of KIF5C(1-560)-Δ6 mutant motors (Figure 1) that results in microtubule destruction, we tested whether expression of other kinesin motors with similar motility properties results in alteration of the microtubule network in COS-7 cells. To do this, we expressed the constitutively-active kinesin-3 motor KIF1A(1-393) which has comparable single-motor motility properties (fast and superprocessive motility, rapid detachment from the microtubule under low forces) [Table 1, [23, 34-36]]. However, expression of KIF1A(1-393) did not lead to the destruction of the microtubule network (Figure 2C). In these cells, the microtubule network was not fragmented (Figure 2D) but rather, expression of KIF1A(1-393) appeared to facilitate microtubule polymerization and/or stabilization as the total length increased and the porosity decreased (Figure 2E,F). Collectively, these results suggest that destruction of the microtubule network in cells is not solely a result of the enhanced motility properties of Δ6 mutant motors, as the microtubule network is not destroyed in cells expressing superprocessive KIF1A motors.

**Table 1:**
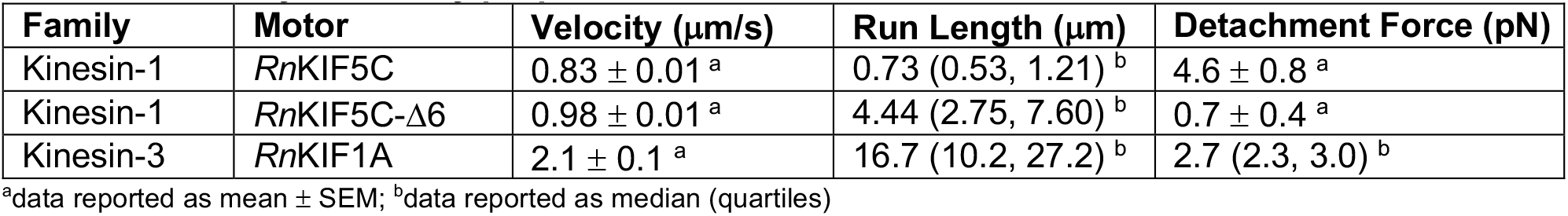
Summary of motility properties of kinesin-1 and kinesin-3 motors.

To rule out the possibility that the effects of the KIF5C(1-560)-Δ6 mutant motor are restricted to COS-7 cells, we expressed KIF5C(1-560)-Δ6 in human hTERT-RPE, MRC-5, U-2-OS cells and in mouse NIH-3T3 cells. In each case, expression of Δ6 motors led to the fragmentation and/or loss of the microtubule network (Figure S3), confirming that this observation is not a cell line-specific artifact. Furthermore, microtubule destruction required processive motility of KIF5C(1-560)-Δ6 as expression of a monomeric version [KIF5C(1-339)-Δ6] did not alter the organization or morphology of the microtubule network (Figure S4).

### KIF5C(1-560)-Δ6 activity leads to buckling, knotting, and breakage of microtubules

We used live-cell imaging to uncover the events leading to destruction of the microtubule network in cells expressing KIF5C(1-560)-Δ6 motors. We imaged microtubule dynamics at the periphery of COS-7 cells using EGFP-α-tubulin and TIRF microscopy. The expressed KIF5C(1-560)-WT, KIF5C(1-560)-Δ6, and KIF1A(1-393) motors were tagged with a Halo tag and labeled with JF552-Halo ligand.

Microtubules in control cells and in cells expressing KIF5C(1-560)-WT motors underwent periods of growth and then paused at the cell periphery before undergoing catastrophe and shrinkage. In these cells, the majority of pauses lasted <50 sec (Figure 3B) and although the paused microtubules often underwent localized buckling in response to compressive forces (25% of microtubules in untransfected cells and 45% in KIF5C(1-560)-WT cells; Figure 3A), the pausing and buckling rarely led to microtubule breakage. In contrast, expression of KIF5C(1-560)-Δ6 mutant motors or KIF1A(1-393) motors resulted in a dramatic increase in the time of pausing at the cell periphery such that the majority of microtubule pauses lasted >150 sec (Figure 3C,D). Furthermore, the paused microtubules showed an increased tendency to buckle (53% of microtubules in KIF5C(1-560)-Δ6 cells and 59% in KIF1A(1-393) cells; Figure 3A). Together, the increased pause time and buckling resulted in highly curled microtubules that tend to loop back towards the center of the cell and become knotted with one another (Figure 2A, arrowheads; Movies 1 and 2).

**Figure 3.**
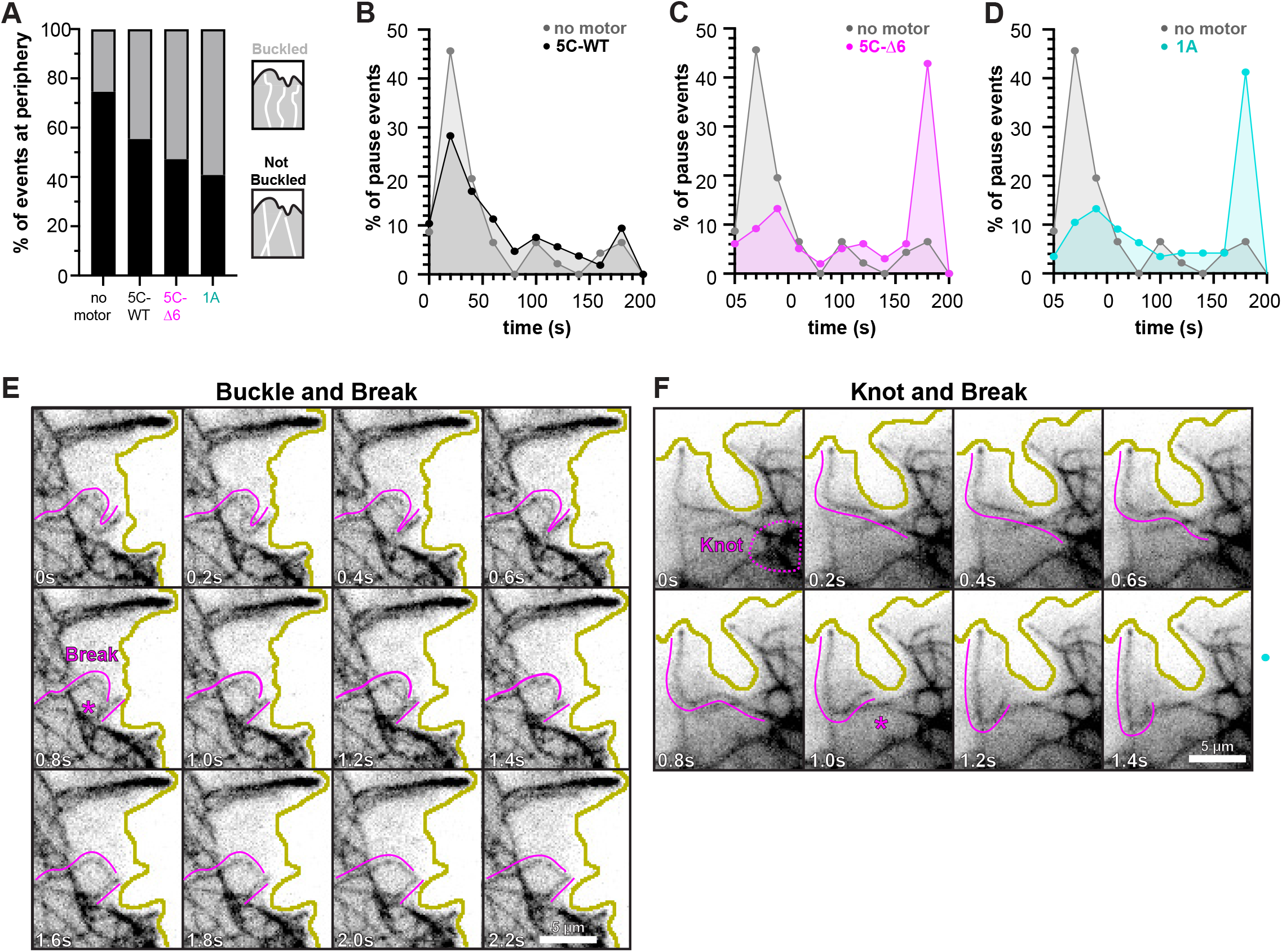
Expression of KIF5C(1-560)-Δ6 leads to buckling, knotting, and breakage of microtubules in cells. (A-D) Quantification of microtubule buckling and pausing at the cell periphery. Microtubule growth events at the cell periphery were scored from movies of cells expressing KIF5C(1-560)-WT, KIF5C(1-560)-Δ6, or KIF1A(1-393) motors. (A) Microtubules that grew to the cell periphery were scored as not buckling (black) or buckling (grey) before catastrophe. The data for each construct are summarized as a stacked bar plot. (B-D) The amount of time microtubules paused at the cell periphery (regardless of buckling) before undergoing catastrophe. The frequency of microtubule pause times for each construct is plotted as a line plot. Gray, untransfected cells; black, KIF5C(1-560)-WT; magenta, KIF5C(1-560)-Δ6; cyan, KIF1A(1-393). For each construct, N > 166 microtubule growth events from 7-12 cells were scored. (E-F) Representative images of microtubule destruction in cells expressing KIF5C(1-560)-Δ6 motors. Magenta traces highlight the microtubule that underwent breakage and the asterisk indicates the time frame when the breakage occurred. Yellow lines indicate the periphery of the cell. Scale bars, 5 μm. (E) A microtubule buckles at the cell periphery and then breaks within the buckled region. (F) A microtubule extends from a region of knotted microtubules (dotted circle) and then breaks near the knot while its plus end remains paused at the cell periphery.

Expression of both KIF5C(1-560)-Δ6 and KIF1A(1-393) motors resulted in increased microtubule bending, looping, and buckling, however, only microtubules in cells expressing the KIF5C(1-560)-Δ6 motors were observed to break into fragments. Microtubule breakage was observed to occur after persistent bending and buckling of a microtubule at the cell periphery (Figure 3E) as well as at locations of dynamic, “knotted” microtubules (Figure 3F). The fragments released after microtubule breakage often underwent depolymerization (Figure S5) and the cumulative effect of microtubule fragmentation and depolymerization was a loss of microtubules at the periphery of the cell (Figure 2, Figure S5, Movie 3). Overall, these results suggest that the activity of KIF5C(1-560)-Δ6 mutant motors leads to bending, buckling, knotting, and breakage of microtubules, a phenomenon most noticeable at the cell periphery, and the eventual destruction of the microtubule network in cells.

### Microtubule acetylation is not sufficient to protect microtubules from KIF5C(1-560)-Δ6 destruction

Recent work has demonstrated that acetylation of α-tubulin at Lysine 40 (αTub-K40ac) weakens lateral interactions between tubulin subunits within the microtubule lattice, thereby increasing microtubule flexibility and the ability to withstand breakage caused by mechanical stress [37-39]. We thus tested whether increasing αTub-K40ac can protect microtubules from destruction caused by the activity of KIF5C(1-560)-Δ6 mutant motors.

We co-transfected COS-7 cells with plasmids for expression of α-tubulin acetyltransferase (αTAT1) tagged with mCit and either KIF5C(1-560)-WT or KIF5C(1-560)-Δ6 mutant motors tagged with Halo-FLAG and labeled with JF552-Halo ligand. Expression of αTAT1 resulted in a dramatic increase in the levels of αTub-K40ac (Figure S6; Figure 4D), as expected from previous work [40, 41]. However, the increased αTub-K40ac provided only minimal protection from breakage caused by KIF5C(1-560)-Δ6 motors as cells expressing the mutant motor still displayed a high number of microtubule fragments in the presence of αTAT1 (Figure 4C,E; Figure S6B), although fewer fragments than cells expressing KIF5C(1-560)-Δ6 in the absence of αTAT1 (Figure 4B,E). Cells expressing KIF5C(1-560)-Δ6 motors in the presence and in the absence of αTAT1 also had similar total microtubule lengths and microtubule density at the cell periphery (Figure 4F,G).

**Figure 4.**
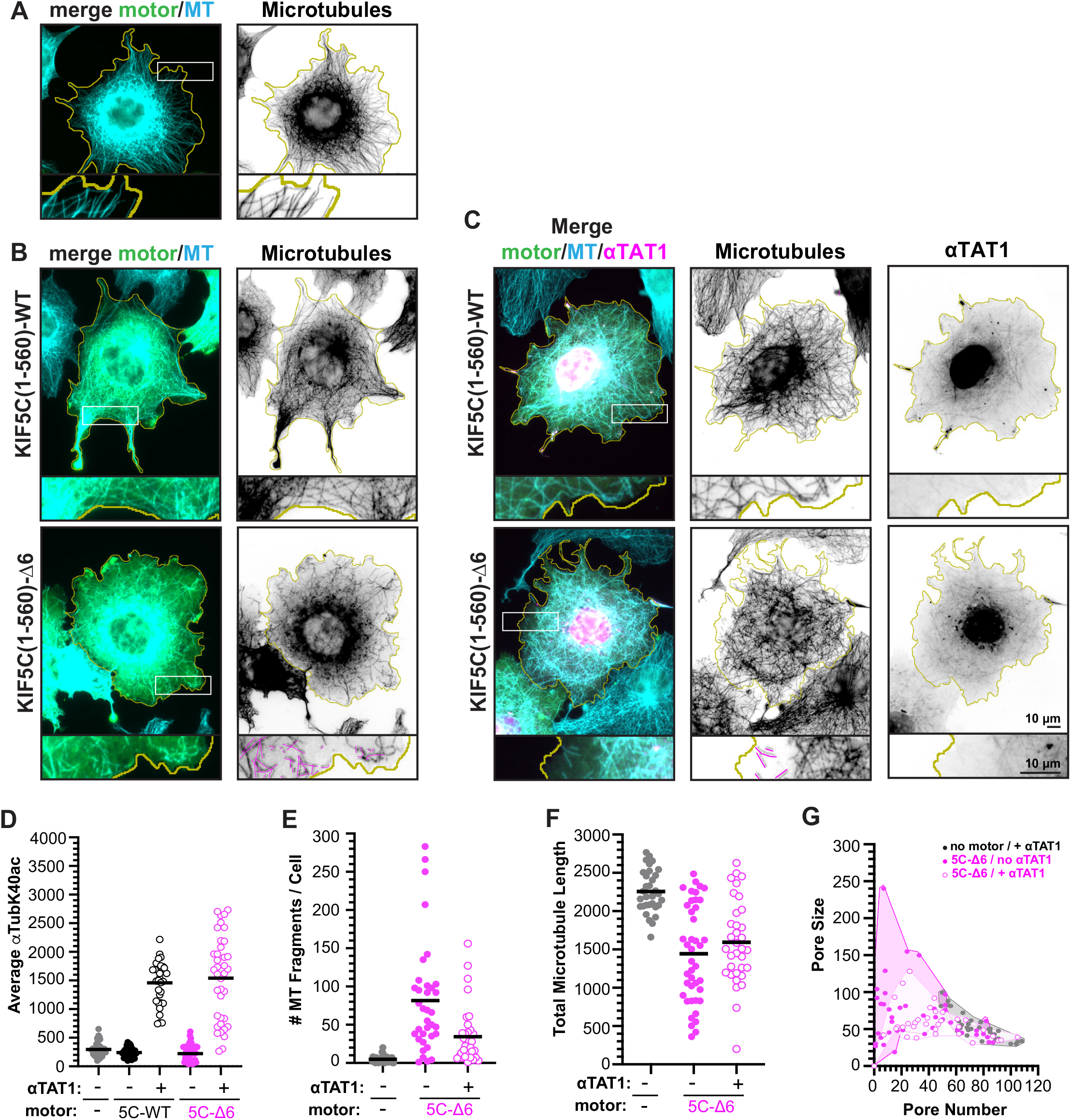
Expression of αTAT1 drives increased microtubule acetylation but does not protect the microtubule network from KIF5C(1-560)-Δ6 destruction. (A-C) Representative images of the microtubule (MT) network in cells expressing (A) no motor or αTAT1, (B) KIF5C(1-560)-WT or KIF5C(1-560)-Δ6 but no αTAT1, or (C) KIF5C(1-560)-WT or KIF5C(1-560)-Δ6 together with αTAT1. The cells were fixed and stained with antibodies against total α-tubulin (microtubules) or αTub-K40ac (not shown, see Figure S6). Yellow lines indicate the periphery of each cell; white boxes indicate regions shown below each image at higher magnification; magenta traces indicate microtubule fragments. Scale bars, 10 μm. (D) Quantification of microtubule acetylation. The average amount of microtubule acetylation was measured for each cell and plotted as a dot plot. N > 46 cells across two independent trials. (E-G) Quantification of changes in the microtubule network. Each dot represents one cell [gray, untransfected cells; black, KIF5C(1-560)-WT; magenta, KIF5C(1-560)-Δ6]. (E) Number of microtubule fragments per cell. (D) Total length of microtubules in a 100×100 pixel box at the cell periphery. (G) Density of microtubules in a 100×100 pixel box at the cell periphery as described by the number and size of the pores in the microtubule network.

As an alternative strategy to assess the impact of αTub-K40ac on motor-driven microtubule destruction, we treated cells with the deacetylase inhibitor trichostatin A (TSA) to block the activity of the α-tubulin deacetylase HDAC6 [42, 43]. Cells treated with TSA had a substantial increase in αTub-K40ac levels (Figure S7), however, expression of KIF5C(1-560)-Δ6 mutant motors still caused destruction of the microtubule network (Figure S7C). We thus conclude that α-tubulin acetylation is not sufficient to protect microtubules from damage and breakage caused by the motility of KIF5C(1-560)-Δ6 motors.

### KIF5C(1-560)-Δ6 motors fail to promote rescue events in microtubule dynamics assays

To gain an understanding of how KIF5C(1-560)-Δ6 impacts microtubules and leads to their destabilization and/or destruction, we turned to *in vitro* assays. We started with a microtubule dynamics assay to explore how the activity of KIF5C(1-560)-WT and KIF5C(1-560)-Δ6 motors leads to the destruction of growing microtubules. Microtubules were grown from GMPCPP-tubulin seeds in presence of soluble GTP-tubulin and either 3xmCit-tagged KIF5C(1-560)-WT or KIF5C(1-560)-Δ6 motors (Figure 5A). In the absence of motor, classic microtubule dynamics of growth, catastrophe, and shrinkage were observed (Figure 5B). After catastrophe, the shrinking microtubules rarely underwent rescue events (resumption of microtubule growth) but rather depolymerized all the way back to the GMPCPP-seed (Figure 5C).

**Figure 5.**
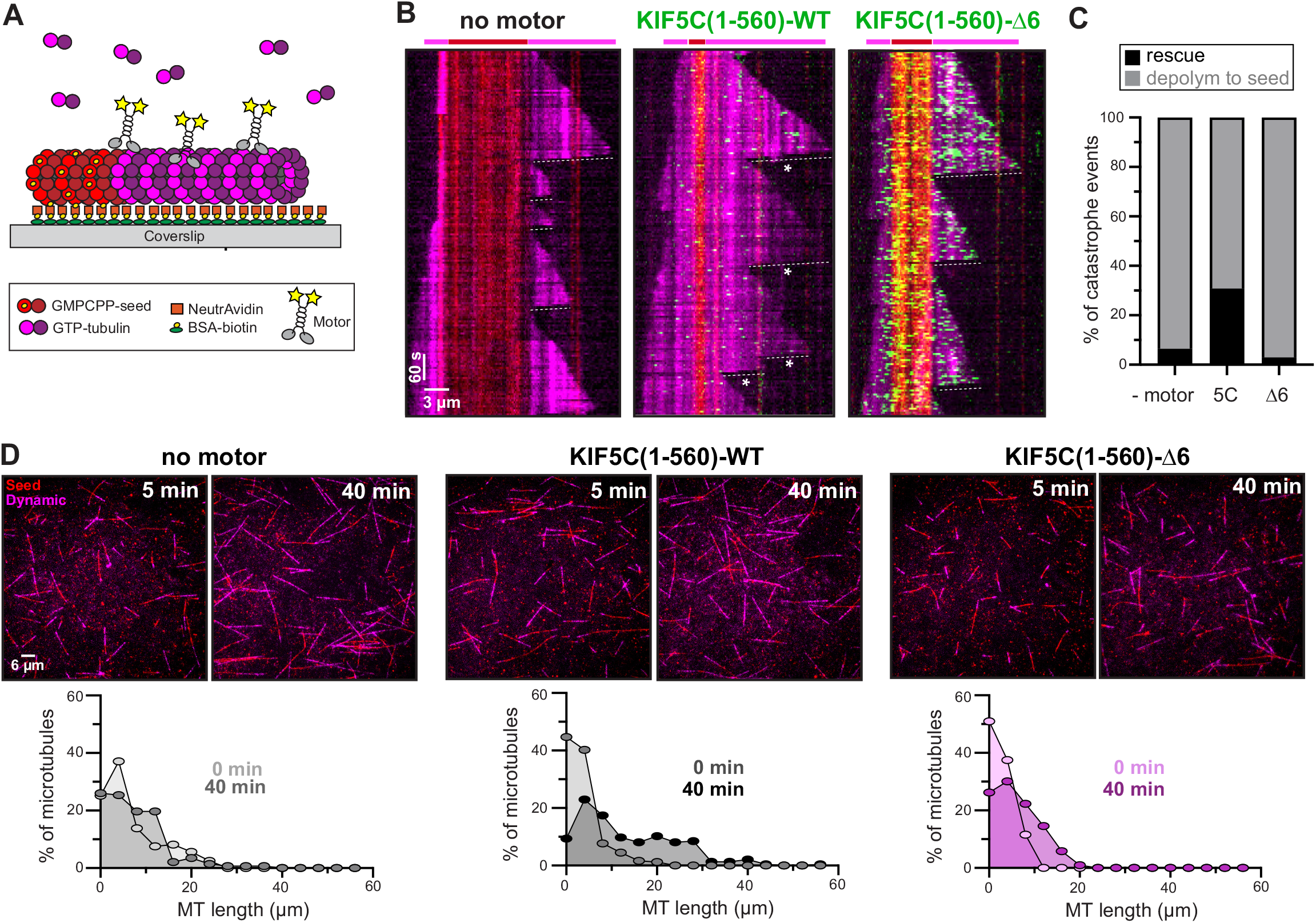
KIF5C(1-560)-WT but not KIF5C(1-560)-Δ6 promotes microtubule rescue and overall microtubule growth. (A) Schematic of microtubule dynamics assay. Microtubules were polymerized from biotinylated GMPCPP-tubulin seeds in the presence of 10.7 μM tubulin and in the absence of motor or presence of 36 nM 3xmCit-tagged KIF5C(1-560)-WT or KIF5C(1-560)-Δ6 motors. (B) Representative kymographs. Red, biotinylated GMPCPP microtubule seeds; magenta, dynamic microtubules; green, KIF5C motors. Time is displayed on the y-axis (scale bar, 60 seconds) and distance on the x-axis (scale bar, 3 μm). Dotted lines indicate catastrophe and depolymerization events; asterisks mark rescue events. (C) Quantification of the frequency of microtubule rescue. From movies of dynamic microtubules, events were scored as a catastrophe followed by depolymerization to the GMPCPP-seed (gray) or catastrophe followed by a rescue event and new microtubule growth before reaching the GMPCPP-seed (black). N > 170 microtubules for each condition across two independent trials. (D) Representative images showing the overall length of the dynamic microtubules 5 min or 40 min after microtubule growth. Red, biotinylated GMPCPP microtubule seeds; magenta, dynamic microtubules. Scale bar, 6 μm. Lower graphs display quantification of the overall length of dynamic microtubules at the beginning (0 min) and end (40 min) of the assay. N > 102 microtubules across two independent trials.

KIF5C(1-560)-WT motors underwent motility along both the GMPCPP-seed and GDP-microtubule lattice and their activity resulted in a dramatic increase in the number of rescue events (Figure 5B,C) and a dramatic increase in the overall length of microtubules in the chamber over the course of the imaging time (Figure 5D, middle panel). KIF5C(1-560)-Δ6 motors were also observed to walk along microtubules (Figure 5B), albeit with increased speed, processivity, and landing rate, yet the activity of the mutant motors did not lead to an increase in the number of rescue events (Figure 5C) nor an increase in the overall length of microtubules (Figure 5D, right panels). Thus, while KIF5C(1-560)-WT motors are able to promote microtubule polymerization by inducing rescue events, KIF5C(1-560)-Δ6 motors fail to promote rescue events.

### KIF5C(1-560)-Δ6 causes more damage to the microtubule lattice than the WT protein

We considered several possible explanations for why KIF5C(1-560)-Δ6 motors are unable to promote rescue events for dynamic microtubules. One possibility is that the weak force output of KIF5C(1-560)-Δ6 motors (Figure 1) results in little damage to the microtubule and thus only rare tubulin repair events that can trigger rescue events. An alternative possibility is that WT and Δ6 motors induce equivalent amounts of damage to the lattice but the defects induced by KIF5C(1-560)-Δ6 motors cannot be repaired simply by the addition of soluble tubulin. A third possibility is that KIF5C(1-560)-Δ6 motors generate more damage than the WT motor such that the repair events are insufficient to maintain microtubule stability.

To test these possibilities, we performed microtubule repair assays in which single kinesin motors walk along immobilized microtubules in the presence of soluble tubulin [17, 44]; the incorporation of soluble tubulin into the microtubule lattice is measured as an indication of the amount of microtubule damage and repair. As motor-induced damage was observed to occur along GDP-tubulin microtubules, but not GMPCPP- or taxol-microtubules [17], we generated GDP-lattice microtubules by growing GTP-tubulin microtubules from GMPCPP-seeds and capping them with GMPCPP-tubulin (Figure 6A). In the absence of motor, GMPCPP-seeded and -capped microtubules are relatively stable and show little to no incorporation of soluble tubulin along the microtubule lattice (Figure 6B). We thus repeated the experiments in the presence of KIF5C(1-560)-WT motors or KIF5C(1-560)-Δ6 motors. As KIF5C(1-560)-Δ6 motors show increased processivity and landing rate on taxol-stabilized and GDP-microtubules (Figures 1,5), we added 3x more KIF5C(1-560)-WT motors to ensure equivalent activity of WT and mutant motors in the assay. In the presence of KIF5C(1-560)-WT motors, small regions of tubulin incorporation into the GDP-microtubule lattice could be detected (Figure 6B). The activity of KIF5C(1-560)-Δ6 motors also resulted in an increase in the incorporation of soluble tubulin into the GDP-microtubule lattice (Figure 6B) but the repair sites induced by the activity of KIF5C(1-560)-Δ6 motors were significantly longer than those caused by the WT motor (Figure 6C). These results indicate that both KIF5C(1-560)-WT and KIF5C(1-560)-Δ6 motors generate defects in the microtubule lattice that can be repaired by the incorporation of soluble tubulin but that the KIF5C(1-560)-Δ6 motor induces larger damage sites that are likely insufficient to maintain microtubule integrity.

**Figure 6.**
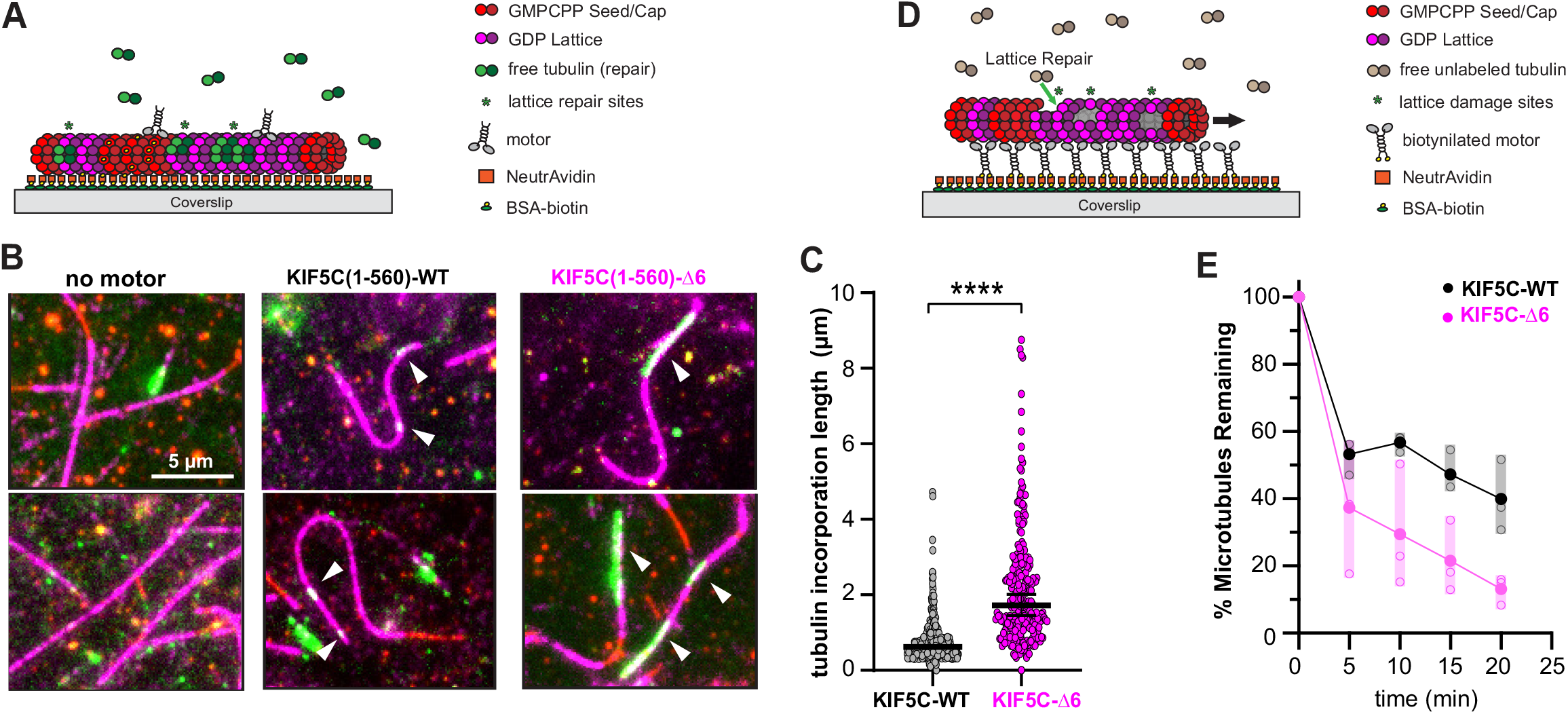
KIF5C(1-560)-Δ6 induces more microtubule damage than KIF5C(1-560)-WT. (A-C) Microtubule repair assay. (A) Schematic of assay. GDP-tubulin microtubules (GMPCPP-seeds and GMP-CPP-caps) were attached to a coverslip. Purified Halo-Flag-tagged KIF5C(1-560)-WT (18 nM) or KIF5C(1-560)-Δ6 (6 nM) motors were added to the flow chamber in the presence of 10 μM 488nm-labeled soluble tubulin. Static images were obtained after 7 min of free tubulin incorporation into motor-driven damage sites. (B) Representative images. Green, microtubule repair sites; red, GMPCPP-seeds; magenta, GDP-tubulin lattice. Arrowheads indicate microtubule repair sites. Scale bar, 5 μm. (C) Quantification of the length of microtubule repair sites. N=375 [KIF5C(1-560)-WT] and N=246 [KIF5C(1-560)-Δ6] repair sites across three independent experiments. ****, p<0.0001 (two-tailed t test). (D,E) Microtubule destruction assay. (D) Schematic of assay. 75 nM of biotinylated KIF5C(1-560)-WT or KIF5C(1-560)-Δ6 motors drive the gliding of GDP-tubulin microtubules (GMPCPP-seeds and GMPCPP-caps) in the presence of 7 μM soluble unlabeled tubulin. (E) Quantification of motor-driven microtubule destruction over time. The total length of microtubules was measured at the indicated time points and the percentage of microtubules remaining in the chamber at the indicated time points was calculated and plotted as a dot plot. Black, KIF5C(1-560)-WT; magenta, KIF5C(1-560)-Δ6. Solid dots indicate the average loss of microtubule length across three independent trials, open dots indicate loss of microtubule length for an individual trial.

To extend these results, we carried out microtubule destruction assays [7, 9-11, 17] in which anchored motors cause stress on the lattice and microtubule destruction even in the presence of soluble tubulin. For these assays, KIF5C(1-560)-WT and KIF5C(1-560)-Δ6 motors were tagged with an Avitag and biotinylated via co-expression with HA-BirA. Motors were adhered to the surface of neutravidin-coated coverslips and then GDP-microtubules stabilized with GMPCPP seeds and caps were added in the presence of soluble tubulin (Figure 6D). While the activity of both KIF5C(1-560)-WT and KIF5C(1-560)-Δ6 motors resulted in the gradual destruction of microtubules over time (Figure 6E), the destruction caused by KIF5C(1-560)-Δ6 motors occurred at a faster rate and to a much greater extent than the damage caused by the activity of WT motors (Figure 6E). These results support the hypothesis that the KIF5C(1-560)-Δ6 motors generate more damage to the microtubule lattice than KIF5C(1-560)-WT motors and that the resulting microtubules are more sensitive to mechanical stress.

Finally, to verify that KIF5C(1-560)-Δ6 causes more damage to the microtubule lattice than KIF5C(1-560)-WT motors, we incubated motors and microtubules on grids for 5 min in the absence of soluble tubulin. After blotting and washing, the microtubules were visualized by negative stain electron microscopy. In the absence of motor, long straight microtubules were observed regardless of polymerization condition (taxol, GMPCPP-tubulin, or GDP-tubulin) (Figures 7). In the presence of KIF5C(1-560)-WT motors, defects and breaks in the lattice were occasionally observed whereas in the presence of KIF5C(1-560)-Δ6 motors, nearly every microtubule showed some type of defect or break in the lattice (Figures 7 and S8). Together, these results support the hypothesis that KIF5C(1-560)-Δ6 motors generate more damage to the microtubule lattice and that the damage makes microtubules susceptible to mechanical stress and breakage.

**Figure 7.**
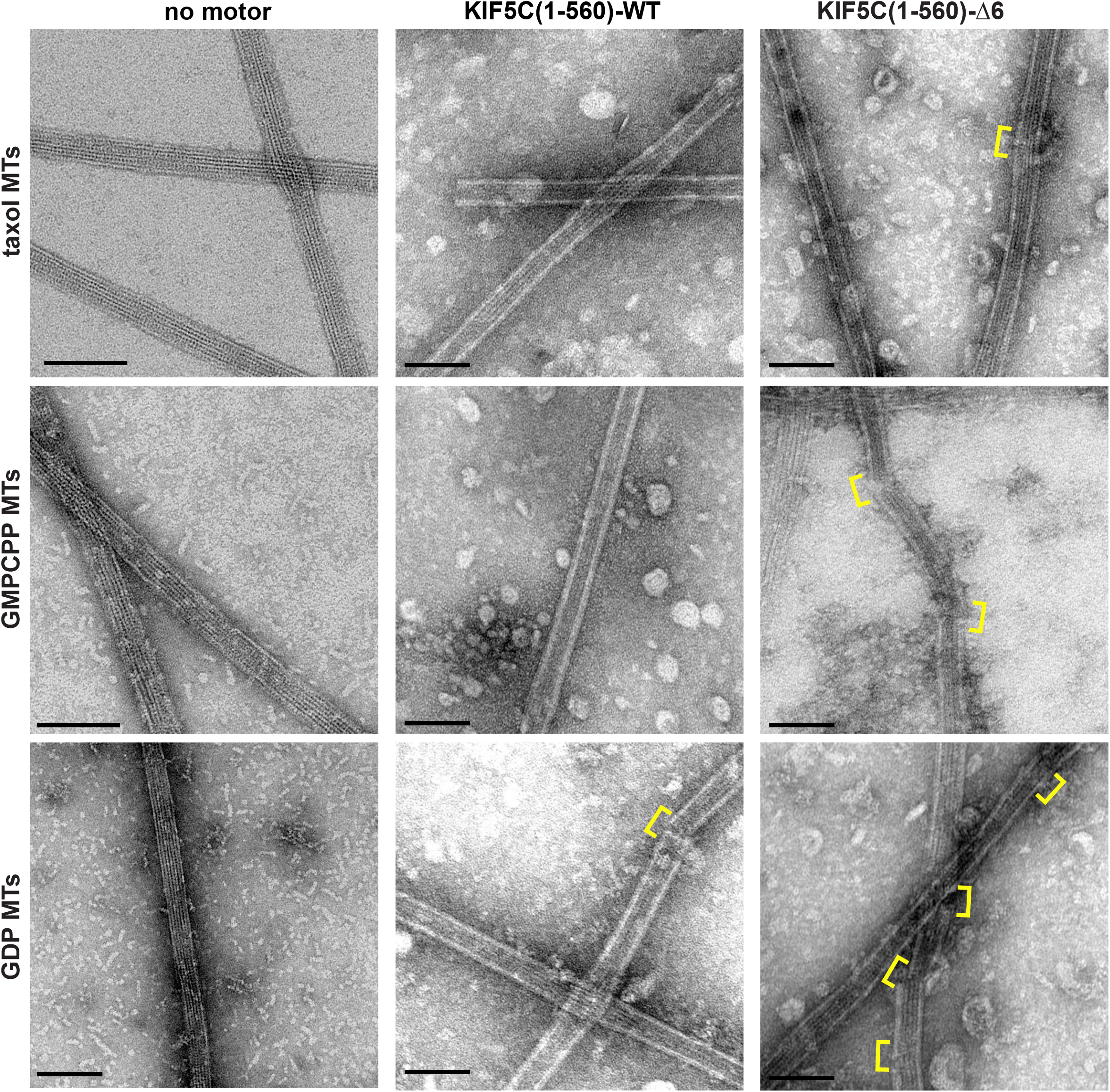
KIF5C(1-560)-Δ6 induces more microtubule damage than KIF5C(1-560)-WT. Taxol-stabilized microtubules (taxol MTs), GMPCPP-tubulin microtubules (GMPCPP MTs), or GDP-tubulin microtubules (GDP MTs) were incubated on grids with no motor or with purified Halo-Flag-tagged KIF5C(1-560)-WT or KIF5C(1-560)-Δ6 motors for 5 min before blotting and staining with uranyl formate. Yellow brackets indicate sites of lattice damage. Scale bars, 100 nm.

## Discussion

### The length of the coverstrand is optimized for kinesin-1 motility and force generation

The mechanistic basis of ATP-dependent force generation by motor proteins continues to be of great interest [4]. For kinesin-1 proteins, zippering of the coverstrand (β0) with the neck linker (β9-β10) to form a two-stranded CNB is a critical first step in generation of a power stroke [21, 31]. CNB formation has been also observed structurally for members of the kinesin-2, kinesin-3, kinesin-5 and kinesin-6 families [27, 45-50] and is critical for kinesin-3 force generation [23].

For CNB formation, variations in both the sequence and length of the coverstrand are thought to dictate kinesin-specific force and motility properties [27]. Previous investigations of how the length of the coverstrand impacts force generation utilized deletions of the entire coverstrand segment that abrogated force output [22, 51]. Here we show that even a partial truncation of the coverstrand results in a kinesin-1 motor that is severely compromised in force generation as KIF5C(1-560)-Δ6 motors detached from the microtubule at low opposing forces (<1 pN, Figure 1). Thus, shortening of the coverstrand had the same effect on kinesin-1 force generation as did single mutations [22, 24].

Shortening of the coverstrand also resulted in kinesin-1 motors with enhanced motility properties (velocity, processivity, landing rate) under unloaded conditions (Figure 1), likely due to allosteric effects of NL docking on the microtubule- and nucleotide-binding regions of the motor domain. While point mutations in the coverstrand of rat and *Drosophila* kinesin-1 motors also resulted in enhanced motility properties [22, 24], deletion of the entire coverstrand of *Drosophila* kinesin-1 had the opposite effect of reduced motility properties under unloaded conditions [22]. Thus, a weakening of CNB formation via point mutation or partial deletion enables the motor to move with greater speed and processivity under unloaded conditions but a complete loss of CNB formation impairs the motor’s ability to efficiently undergo processive motility. These findings support the hypothesis that the sequence and length of the coverstrand have been optimized in order to balance the motility and force generation properties of processive kinesins [22, 24].

### Kinesin-1 motor activity causes microtubule fragmentation in cells

We demonstrate that kinesin-1 motility causes microtubule breakage and fragmentation in cultured cells. For the WT kinesin-1 KIF5C(1-560), microtubule fragmentation was observed only when the motor was expressed at high levels. In contrast, even low levels of expression of the KIF5C(1-560)-Δ6 variant caused microtubule breakage and fragmentation (Figure 2, Figure S3). We also observed microtubule destruction in cells upon expression of other KIF5C variants with truncation (Δ3, Δ7, Δ8, Δ9) or mutation [24] of the coverstrand. These results extend previous reports demonstrating that kinesins and dyneins can cause microtubule destruction in reconstituted systems [17, 44] to show that motor-induced damage occurs in cells. These findings emphasize that while cells are able to repair motor-induced damage under normal conditions, and even upon overexpression of active kinesin-1 motors, the excessive ability of the Δ6 variant to damage microtubules overwhelms the cells’ ability to repair and/or resolve lattice defects, resulting in broken and fragmented microtubules.

Not all kinesins cause microtubule breakage and fragmentation as expression of the kinesin-3 motor KIF1A resulted in the formation of highly curved and looped microtubules but not their fragmentation. We speculate that the ability of kinesin-1 KIF5C motors, but not kinesin-3 KIF1A motors, to induce defects in the microtubule lattice is due to differences in the chemomechanical cycles of these motors. Kinesin-1 motors spend a majority of their mechanochemical cycle with both heads strongly-bound to the microtubule (two-head-bound state), and their residence time in this state increases in response to force [36, 52-55]. In contrast, the kinesin-3 motor KIF1A spends the majority of its mechanochemical cycle in a one-head bound state, tethered to the microtubule via electrostatic interactions that enable superprocessive motility under unloaded conditions but detachment from the microtubule at relatively low resistive forces [23, 36, 56-60]. Thus, sequence changes that optimized kinesin-1’s mechanochemistry for high-load transport appear to have rendered it susceptible to destruction of the microtubule track that it walks along.

Using live-cell imaging, we observed that both kinesin-1 KIF5C(1-560)-Δ6 and kinesin-3 KIF1A(1-393) motors accumulate at the plus ends of the microtubules in the cell periphery. Motor accumulation results in an increased time of microtubule pausing at the cell periphery and this, in turn, results in microtubule bending, looping, and buckling (Figure 3). Microtubule bending and buckling occurs in response to compressive loads but does not normally lead to microtubule breakage and fragmentation [61-68]. We suggest that the excessive damage caused by the KIF5C(1-560)-Δ6 variant results in microtubules that break under compressive loads.

### Motor-induced damage makes microtubules sensitive to mechanical stress

Using *in vitro* reconstitution assays, we demonstrate that the activity of KIF5C(1-560)-WT and KIF5C(1-560)-Δ6 motors results in an increase in the frequency of lattice repair sites as compared to the no motor condition (Figure 6). We also demonstrate that the activity of KIF5C(1-560)-Δ6 motors results in an increase in the size of the lattice repair sites compared to the KIF5C(1-560)-WT motor (Figure 6). Together with recent reports [17, 44], our work suggests that while stepping along the microtubule lattice, kinesin-1 motor proteins induce defects in the microtubule lattice that can be repaired by the incorporation of soluble tubulin.

Lattice repair in response to KIF5C(1-560)-WT activity led to an increase in microtubule rescue events and overall microtubule growth (Figure 5), as also reported by [44]. However, the excessive damage caused by KIF5C(1-560)-Δ6 motors and insufficient repair of those defects resulted in a lack of rescue events (Figure 5). The excessive damage caused by KIF5C(1-560)-Δ6 motors also resulted in fragmentation and destruction of microtubules under mechanical stress: in microtubule gliding assays (Figure 6), in motility assays on EM grids (Figure 7), and when bent and buckled in cells (Figures 2,3).

The ability of microtubules to bear significant compressive loads has been documented as, for example, hundreds of pN of force were needed to rupture microtubules in *in vitro* assays [69, 70] and microtubules have been documented to bear similarly-large compressive forces in cells [61]. Thus, how processive motors generating ∼5 pN of force could cause microtubules to break and fragment was unclear. Our results indicate that kinesin-1 motility leads to the loss of tubulin subunits within the microtubule lattice which reduces the microtubule’s flexural rigidity. For KIF5C(1-560)-WT, this softening effect allows the microtubules to bend, buckle, and bear significant compressive forces, as has been noted in cells expressing members of the MAP65/PRC1/Ase1 family [71]. In the case of KIF5C(1-560)-Δ6, however, the excessive defects in the microtubule lattice result in microtubules that are “too soft” and undergo fragmentation when experiencing mechanical stress in both *in vitro* assays and in cells.

### Repair of motor-induced damage in cells

An important finding of our work is that repair of lattice defects is essential to prevent microtubule breakage and fragmentation in cells. Indeed, lattice repair could be part of a positive feedback loop that results in strong and stable microtubule tracks [20]. Defects along cellular microtubules can be observed by cryoEM [63, 72-74] and their repair is likely to be particularly important in neurons where the microtubules span the length of the axon and in cardiomyocytes where microtubules bear significant compressive loads [75]. An important corollary is that cells have a limited capacity to repair or resolve lattice defects and conditions that exceed this capacity, such as expression of highly destructive KIF5C(1-560)-Δ6 motors, result in microtubule breakage and fragmentation. Our identification of Δ6 as a kinesin motor that enhances microtubule damage in cells enables future work exploring whether lattice repair mechanisms become limiting for kinesin and/or tubulin mutations that lead to neurodegenerative diseases.

Our identification of Δ6 as a kinesin motor that enhances microtubule damage in cells also enables future work to decipher the roles of specific cellular factors in microtubule repair. That is, while the incorporation of soluble tubulin is sufficient to reduce microtubule damage in *in vitro* assays, additional factors may contribute to recognition and repair of microtubule damage in cells. These factors include the end binding (EB) proteins which accumulate at damage sites in vitro [75, 76] and can accumulate as islands on the microtubule lattice in cells [77], CLIP-170 (cytoplasmic linker protein of 170 kDa) and related proteins that promote microtubule rescues [78-80], and CLASPs (cytoplasmic linker associated proteins) which can bind tubulin heterodimers and oligomers as well as microtubules [81-83] and have been proposed to contribute to microtubule repair [84].

## Materials and Methods

### Plasmids and Adenoviral vectors

A truncated, constitutively active kinesin-1 [rat KIF5C(1-560)] was used. Coverstrand truncation mutants were generated by PCR and all plasmids were verified by DNA sequencing. The truncated, constitutively active version of the kinesin-3 motor KIF1A contains the first 393 amino acids of rat KIF1A, which includes the neck coil sequence for dimerization, followed by a Leucine zipper (LZ) sequence to maintain the dimer state [35]. Motors were tagged with three tandem monomeric Citrine fluorescent proteins (3xmCit) or HALO-FLAG tags for single-molecule imaging assays, a FLAG tag for optical trapping assays, monomeric NeonGreen (mNG)-FRB for Golgi dispersion assays, or were biotinylated via an AviTag (aa sequence GLNDIFEAQKIEWHE) and co-expression with HA-BirA for microtubule gliding assays. The mouse αTAT1 coding sequence (NP_001136216) was cloned into the vector pmCit-C1. The Golgi-targeting GMAP-mRFP-FKBP construct contains the Golgi-targeting sequence of HsGMAP210 (amino acids 1757-1838, NP_004230) [85]. Constructs coding for FRB (DmrA) and FKBP (DmrC) sequences were obtained from ARIAD Pharmaceuticals and are now available from Takara Bio Inc. Plasmids encoding monomeric NeonGreen were obtained from Allele Biotechnology. The adenovirus plasmid encoding EGFP-tubulin (pShuttle-EGFP-tubulin) was a gift from Torsten Wittmann (Addgene plasmid #24327, RRID:Addgene_24327, [86]) and adenovirus was produced by the University of Michigan Vector Core.

### Cell culture, transfection, and lysate preparation

COS-7 (African green monkey kidney fibroblasts, American Type Culture Collection, RRID: CVCL_0224) were grown at 37°C with 5% (vol/vol) CO_2_ in Dulbecco’s Modified Eagle Medium (Gibco) supplemented with 10% (vol/vol) Fetal Clone III (HyClone) and 2 mM GlutaMAX (L-alanyl-L-glutamine dipeptide in 0.85% NaCl, Gibco). Cells are checked annually for mycoplasma contamination and were authenticated through mass spectrometry (the protein sequences exactly match those in the African green monkey genome). 24 hr after seeding, the cells were transfected with plasmids using TransIT-LT1 transfection reagent (Mirus) and Opti-MEM Reduced Serum Medium (Gibco). Cells were processed 24 hr after transfection. For lysates, the cells were trypsinized and harvested by centrifugation at 3000 × *g* at 4**°**C for 3 min. The pellet was resuspended in cold 1X PBS, centrifuged at 3000 × *g* at 4**°**C for 3 min, and the pellet was resuspended in 50 μL of cold lysis buffer [25 mM HEPES/KOH, 115 mM potassium acetate, 5 mM sodium acetate, 5 mM MgCl_2_, 0.5 mM EGTA, and 1% (vol/vol) Triton X-100, pH 7.4] with 1 mM ATP, 1 mM phenylmethylsulfonyl fluoride, and 1% (vol/vol) protease inhibitor cocktail (P8340, Sigma-Aldrich). Lysates were clarified by centrifugation at 20,000 × *g* at 4**°**C for 10 min and lysates were snap frozen in 5 μL aliquots in liquid nitrogen and stored at −80**°**C.

### Imaging of fixed and live cells

For fixed cells, 24 hr post-transfection, the cells were rinsed with PBS and fixed in 3.7% (vol/vol) paraformaldehyde (ThermoFisher Scientific) in PBS for 10 min at room temperature. Fixed cells were permeabilized in 0.2% Triton X-100 in PBS for 5 min and blocked with 0.2% fish skin gelatin in PBS for 5 min. Primary and secondary antibodies were applied in blocking buffer for 1 hr at room temperature in the dark. Primary antibodies: Ms anti-β-tubulin (clone E7, Developmental Studies Hybridoma Bank, 1:2000), Ms anti-αTubK40ac (clone 6-11B-1, Sigma T7451; 1:10,000), Rb anti-giantin (Biolegend #924302, 1:200). Secondary antibodies were purchased from Jackson ImmunoResearch Laboratories and used at 1:500 dilution. Nuclei were stained with 10.9 mM 40,6-diamidino-2-phenylindole (DAPI) and the coverslips were mounted using Prolong Gold (Invitrogen). Images were acquired on an inverted epifluorescence microscope (TE2000E; Nikon) with a 40× 0.75 NA, a 60xv1.40 NA oil-immersion, or a 100× 1.40 NA objective and a CoolSnap HQ camera (Photometrics). The fluorescence images were analyzed using ImageJ (National Institutes of Health).

For live-cell imaging of dynamic microtubules, cells in glass-bottom dishes (Matek) were infected with adenovirus for expression of EGFP-α-tubulin. 24 hr later the cells were transfected and Janelia Fluor 552 (JF552, 50 nM, Janelia Materials) ligand was added to label Halo-tagged motors. 16 h post-transfection, the cells were washed and then incubated in Leibovitz’s l-15 medium (Gibco) and imaged at 37°C in a temperature-controlled and humidified stage-top chamber (Tokai Hit). Live-cell imaging was performed on an inverted TIRF microscope T*i*-E/B (Nikon) equipped with the perfect focus system, a 100× 1.49 NA oil immersion TIRF objective (Nikon), three 20-mW diode lasers (488 nm, 561 nm, and 640 nm), and an electron-multiplying charge-coupled device detector (iXon X3DU897; Andor). The angle of illumination was adjusted for maximum penetration of the evanescent field into the cell. Image acquisition was controlled with Elements software (Nikon). Microtubule growth events at the cell periphery were scored from movies of cells expressing no motor, KIF5C(1-560)-WT, KIF5C(1-560)-Δ6, or KIF1A(1-393) motors. Microtubules that grew to the cell periphery were scored as not buckling or buckling before undergoing a catastrophe and depolymerization towards the cell center.

### Optical trapping assays

Bovine brain tubulin (Cytoskeleton TL238) was reconstituted in 25 mL BRB80 buffer [80 mM PIPES (Sigma P-1851), 1 mM EGTA (Sigma E-4378), 1 mM MgCl_2_ (Mallinckrodt H590), pH adjusted to 6.9 with KOH] supplemented with 1 mM GTP (Cytoskeleton BST06) and kept on ice. 13 mL PEM104 buffer (104 mM PIPES, 1.3 mM EGTA, 6.3 mM MgCl_2_, pH adjusted to 6.9 with KOH) was mixed with 2.2 mL 10 mM GTP, 2.2 mL DMSO, and 4.8 mL of 10 mg/mL tubulin and microtubules were polymerized by incubation for 40 min at 37°C. Subsequently, 2 mL of stabilization solution [STAB: 38.6 mL PEM80, 0.5 mL 100 mM GTP, 4.7 mL 65 g/L NaN3 (Sigma S-8032), 1.2 mL 10 mM Taxol (Cytoskeleton TXD01), 5 mL DMSO (Cytoskeleton)] was added to the stock microtubule solution at room temperature.

Optical trap assays were performed as described previously [24, 28, 29]. 0.44 μm anti-FLAG-coated beads were prepared by crosslinking anti-FLAG antibodies (Thermo Fisher Scientific) to carboxy polystyrene beads (Spherotech) via EDC chemistry. Lysates containing FLAG-tagged motors were diluted in assay buffer [AB: P12 buffer (12 mM PIPES (Sigma P-1851), 1 mM EGTA (Sigma E-4378), 1 mM MgCl2 (Mallinckrodt H590), pH adjusted to 6.9 with KOH), 1 mM DTT (Sigma Aldrich), 20 mM Taxol (Cytoskeleton), 1 mg/mL casein (Blotting-Grade Blocker, Biorad), 1 mM ATP (Sigma Aldrich)] and then incubated with gently sonicated anti-FLAG beads to allow binding for 1 hr at 4**°**C on a rotator in the presence of oxygen scavenging reagents (5 mg/mL b-D-glucose (Sigma Aldrich), 0.25 mg/mL glucose oxidase (Sigma Aldrich), and 0.03 mg/mL catalase (Sigma Aldrich).

A flow cell that holds a volume of ∼15 µL was assembled using a microscope slide, etched coverslips, and double-sided sticky tape. Before assembly, etched coverslips were incubated in a solution of 100 µL poly-l-lysine (PLL, Sigma P8920) in 30 mL ethanol for 15 min. The coverslip was then dried with a filtered air line. After flow cell assembly, microtubules were diluted 150 times from the stock in a solution of PemTax (1 µL 10 mM Taxol in 500 µL P12). The diluted microtubules were added to the flow cell and allowed to adhere to the PLL surface for 10 min. Unbound microtubules were then washed out with 20 µL PemTax. A solution of casein (Blotting-Grade Blocker, Biorad 1706404) diluted in PemTax (1:8 mixture) was then added to the flow cell and allowed to incubate for 10 min to block the remainder of the surface to prevent non-specific binding. After the incubation, the flow cell was washed with 50 µL PemTax and 80 µL assay buffer (AB). 20 µL of the bead/motor incubation was then added to the flow cell.

Optical trapping measurements were obtained using a custom-built instrument with separate trapping and detection systems. The instrument setup and calibration procedures have been described previously (Khalil et al., 2008). Briefly, beads were trapped with a 1,064 nm laser that was coupled to an inverted microscope with a 100x/1.3 NA oil-immersion objective. Bead displacements from the trap center were recorded at 3 kHz and further antialias filtered at 1.5 kHz. To ensure that we were at the single molecule limit for the motility assay, the protein-bead ratio was adjusted such that only 5-10% of trapped beads showing microtubule binding. A motor-coated bead was trapped in solution and subjected to position calibration and trap stiffness Labview routines. Afterward, the bead was brought close to a surface-bound microtubule to allow for binding. Bead position displacement and force generation were measured for single motor-bound beads. Detachment force measurements include motility events where single motors reached a plateau stall before detachment and events where the motor abruptly detached from the microtubule. Detachment forces are plotted as a dot plot where each dot indicates the maximum detachment force of an event and the mean for each construct is indicated by a black horizontal line. Statistical differences between the maximum detachment force of wild type and mutant motors were calculated by using a two-tailed unpaired Student’s *t* test.

### Unloaded, single-molecule motility assays

Microtubules were polymerized (unlabeled and HiLyte-647-labeled porcine brain tubulin, Cytoskeleton Inc #T240 and #TL670M) in BRB80 buffer (80 mM Pipes/KOH pH 6.8, 1 mM MgCl_2_, 1 mM EGTA) supplemented with 2 mM GTP and 2 mM MgCl_2_ and incubated for 60 min at 37°C. Taxol (Cytoskeleton Inc) in prewarmed BRB80 was added to 2 μM and incubated for 60 min. Microtubules were stored in the dark at room temperature for up to 2 weeks. Flow cells were prepared by attaching a #1.5 coverslip (ThermoFisher Scientific) to a glass slide (ThermoFisher Scientific) using double-sided tape. Microtubules were diluted in fresh BRB80 buffer supplemented with 10 μM taxol, infused into flow cells, and incubated for four minutes to allow for nonspecific absorption to the glass. Flow cells were incubated with (i) blocking buffer [30 mg/mL casein in P12 buffer (12 mM Pipes/KOH pH 6.8, 1 mM MgCl_2_, 1 mM EGTA) supplemented with 10 μM taxol] for four minutes and then (ii) motility mixture (0.5–1.0 μL of COS-7 cell lysate, 25 μL P12 buffer, 15 μL blocking buffer, 1 mM ATP, 0.5 μL 100 mM DTT, 0.5 μL of 20 mg/mL glucose oxidase, 0.5 μL of 8 mg/mL catalase, and 0.5 μL 1 M glucose). Flow chambers were sealed with molten paraffin wax and imaged on an inverted Nikon Ti-E/B TIRF microscope with a perfect focus system, a 100 × 1.49 NA oil immersion TIRF objective, three 20 mW diode lasers (488 nm, 561 nm, and 640 nm) and EMCCD camera (iXon^+^ DU879; Andor). Image acquisition was controlled using Nikon Elements software and all assays were performed at room temperature.

Motility data were analyzed by first generating maximum intensity projections to identify microtubule tracks (width = 3 pixels) and then generating kymographs in Fiji/ImageJ (National Institutes of Health). Only motility events that lasted for at least three frames were analyzed. Furthermore, events that ended as a result of a motor reaching the end of a microtubule were included; therefore, the reported run lengths for highly processive motors are likely to be an underestimation. For each motor construct, the velocities and run lengths were binned and a histogram was generated by plotting the number of motility events for each bin. The distributions of motor velocities were fit to a Gaussian cumulative and a student’s t test was used to assess whether velocity distributions were significantly different between motors. The cumulative distribution of WT motor run lengths was fit to an exponential distribution as previously described [24, 58]. However, a fit to an exponential decay function was not an appropriate model to describe the cumulative distributions of the Δ6 mutant motor. Rather, the distribution of the run length was fit to a gamma distribution. The expected mean run length was calculated by multiplying the shape and scale parameters. A Kuskal-Wallis one-way analysis of variance was used to assess whether run length distributions were significantly different between motors.

### Protein expression and purification

COS-7 cells were transfected with plasmids for expression of KIF5C(1-560)-WT-Halo-Flag or KIF5C(1-560)-Δ6-Halo-Flag and the protein was fluorescently labelled by the inclusion of 50 nM JF552 Halo ligand (Tocris Bioscience) in the growth medium. Cells from two 10cm dishes were harvested 24 h after transfection and lysed in 1 ml lysis buffer [25 mM HEPES, 115 mM KOAc, 5 mM NaOAc, 5 mM MgCl_2_, 0.5 mM EGTA, 1% Triton X-100, pH to 7.4 with KOH] supplemented with protease inhibitor cocktail, 1 mM PMSF, 1 mM ATP, and 1 mM DTT. After centrifugation at 16,000xg for 10 min at 4°C, the supernatant was incubated with 50 μl anti-Flag M2 agarose beads (Sigma-Aldrich) with rotation for 1.5 h at 4°C. The beads were washed with wash buffer (150 mM KCl, 20 mM Imidazole pH 7.5, 5 mM MgCl_2_, 1 mM EDTA, 1 mM EGTA) supplemented with protease inhibitor cocktail, 1 mM PMSF, 1 mM DTT, and 3 mM ATP, and washed again with wash buffer supplemented with protease inhibitor cocktail, 1 mM PMSF and 1 mM DTT. The protein was eluted with 80 μl BRB80 buffer (80 mM PIPES/KOH pH6.8, 1 mM MgCl_2_, 1 mM EGTA) supplemented with protease inhibitor cocktail, 1 mM PMSF, 0.5 mM DTT, 0.1 mM ATP, 0.5 mg/ml 3xFlag peptide (Sigma-Aldrich) for 1h. The protein was collected as the supernatant after centrifugation at 1,500xg for 5 min at 4°C. Aliquots were snap-frozen in liquid nitrogen and stored at -80°C.

### Microtubule dynamics assay

Microtubule seeds were prepared by polymerizing 25 μM tubulin (Cytoskeleton Inc) consisting of 6% biotinylated-tubulin (Cytoskeleton Inc) and 6% fluorescent (X-Rhodamine or HiLyte647) tubulin (Cytoskeleton Inc) in the presence of the nonhydrolyzable GTP analogue GMPCPP (Jena Bioscience) in BRB80 buffer and 2.5 mM MgCl_2_ for 35 min at 37 **°**C. The seeds were sedimented by centrifugation at 90,000 rpm for 5 min at 25 **°**C (Beckman Coulter). The microtubule pellet was resuspended in warm BRB80 buffer and microtubule seeds were stored in the dark at room temperature.

A flow chamber (∼10 μl volume) was assembled by attaching a clean #1.5 coverslip (Thermo Fisher Scientific) to a glass slide (Thermo Fisher Scientific) with two stripes of double-sided tape. Microtubule seeds were immobilized by sequential incubation with: (i) 1 mg/ml BSA-biotin (A8549; Sigma-Aldrich), (ii) blocking buffer (1 mg/ml BSA in BRB80), (iii) 0.5 mg/ml NeutrAvidin (31000; Thermo Fisher), (iv) blocking buffer, (v) short GMPCPP-stabilized microtubule seeds, and (vi) blocking buffer. Microtubule growth was initiated by flowing in 10.7 μM tubulin containing 7% Hilyte647–labeled tubulin (Cytoskeleton Inc.) together with 36 nM motor proteins in the reaction buffer (BRB80 with 1 mM GTP, 2.5 mM ATP, 0.1 mg/ml BSA, 1 mg/ml casein, 1 mM MgCl2, 0.1% methylcellulose, and oxygen scavenging mix [1 mM DTT, 10 mM glucose, 0.1 mg/ml glucose oxidase, and 0.08 mg/ml catalase]). The flow cells were sealed with molten paraffin wax and imaged by TIRF microscopy. The temperature was set at 37°C in a temperature-controlled chamber (Tokai Hit) and time-lapse images were acquired in 488-nm, 561-nm, and 640-nm channels at a rate of every 5 s for 15 min. Maximum-intensity projections were generated and kymographs (width = 3 pixels) were generated using Fiji/ImageJ and displayed with time on the x-axis and distance on the y-axis. From kymographs, the total number of growth events resulting in catastrophe were determined and scored as either an event resulting in complete depolymerization to the GMPCPP seed or a rescue event followed by new microtubule growth before reaching the GMPCPP seed. The fraction of catastrophe events resulting in rescue are plotted as a stacked bar plot for N >170 microtubules across two independent experiments.

To quantify overall microtubule growth over the course of imaging, still images were acquired in 488-nm, 561-nm, and 640-nm channels at 5 or 40 minutes. The total length of microtubules in each field of view was measured using Fiji/ImageJ (National Institutes of Health) and the measurements were summed across 4 fields of view for each time point from two independent experiments.

### Microtubule repair assay

Microtubule seeds were attached to the surface of a flow chamber by sequential incubation with: (i) 1 mg/ml BSA-biotin for 3 min, (ii) blocking buffer, (iii) 0.5 mg/ml NeutrAvidin for 3 min, (iv) blocking buffer, (v) GMPCPP-stabilized microtubule seeds (18% x-rhodamine tubulin) for 3 min, and (vi) blocking buffer. Microtubules were polymerized from the seeds by incubating 26 μM tubulin [gift of R. Ohi (University of Michigan) with 12.5% HiLyte647-tubuiln (Cytoskeleton. Inc.)] and 1 mM GTP in imaging buffer [BRB80 buffer supplemented with 0.1% methylcellulose, 1 mg/ml casein, 3 mM MgCl_2,_ 6 mM DTT and oxygen scavenger mix (16 mM glucose, 0.7 mg/ml catalase and 0.3 mg/ml glucose oxidase)] for 15 min. at 37**°**C. Microtubules were capped by incubating with 13 μM unlabeled tubulin and 1 mM GMPCPP in imaging buffer for 5 min at 37**°**C. Wash buffer was flowed in to depolymerize the dynamic tubulin structures grown on the stabilizing GMPCPP cap. Subsequently, a mix containing10 μM HiLyte488-tubulin, 1 mM GTP, 5 mM ATP, and purified motors in imaging buffer was flowed in. To achieve equivalent densities of KIF5C(1-560)-WT and KIF5C(1-560)-Δ6 on the microtubules in this assay, we calculated their relative affinities for GDP microtubules in the microtubule dynamics assay (# motors/μm GDP-microtubule/time frame). As KIF5C(1-560)-Δ6 displayed a 3-fold higher density than KIF5C(1-560)-WT on GDP microtubules, a higher concentration of KIF5C(1-560)-WT-Halo-Flag (18 nM) than KIF5C(1-560)-Δ6-Halo-Flag (6 nM) was incubated with the microtubules in this assay. After incubation of motors with microtubules in the presence of soluble tubulin for 7 min at 37**°**C, the flow chamber was washed with blocking buffer to remove unincorporated HiLyte488-tubulin and unbound motors and then 13 μM unlabeled tubulin was added to prevent microtubule depolymerization. The chamber was sealed with molten paraffin wax, and images were collected on a Nikon Ti-E/B TIRF microscope equipped with a 100X 1.49 N.A. oil immersion TIRF objective (Nikon), three 20 mW diode lasers (488 nm, 561 nm and 640 nm), and EMCCD detector (iXon X3DU897, Andor). Microtubule repair sites were defined as sites of HiLyte488-tubulin incorporation site flanked by HiLyte647-tubulin containing microtubule lattice on both sides. The length of incorporation sites was quantified using Fiji/imageJ2.

### Microtubule destruction assay

Biotinylated GMPCPP-seeds and biotinylated KIF5C(1-560)-Avitag motors were attached to the surface of a flow chamber by sequential incubation (5 min each) with (i) 1 mg/ml BSA-biotin, (ii) wash buffer (1 mg/ml BSA in BRB80), (iii) 0.5 mg/ml NeutrAvidin, (iv) wash buffer, (v) GMPCPP-seeds (6% HiLyte647-tubulin, 6% biotin-tubulin) in BRB80, (vi) wash buffer, (vii) cell lysates containing 75 nM biotinylated motors in BRB80 with 3.5 mM ADP, 35 mM glucose, 0.1 U Hexokinase to prevent motors from binding to soluble tubulin or GMPCPP-seeds, and (viii) wash buffer with 3.5 mM ADP. Next the chamber was placed at 37°C in a temperature-controlled and humidified stage-top incubator (Tokai Hit). GDP-lattice microtubules were assembled in the chamber by polymerization of GTP-tubulin for 30 min [7 uM bovine brain tubulin (gift of R. Ohi, University of Michigan) with 6% HiLyte488-tubulin (Cytoskeleton Inc.) in BRB80 with GTP, methylcellulose, BSA, DTT, and ADP (to prevent motor-tubulin binding) followed by polymerization of GMPCPP-caps for 1.5 min [7 uM bovine brain tubulin (gift of R. Ohi, University of Michigan) with 11% HiLyte647-tubuiln (Cytoskeleton. Inc.) in BRB80 with 2.6 mM GMPCPP, methylcellulose, BSA, and ADP (to prevent motor-tubulin binding). The flow chamber was washed twice with wash buffer (P12 buffer (12 mM Pipes/KOH pH 6.8, 1 mM MgCl_2_, 1 mM EGTA) containing ADP) and images of microtubules in 8 fields of view were obtained (0 min time point). Then Motility Mix [imaging buffer (25 ul P12 buffer, 15 ul P12 block (15 mg/mL BSA in P12), 2 ul of 10 mg/mL casein/P12, and 0.5 ul of the following: 1 M DTT, 100 mM MgCl_2_, 1 M glucose, 8 mg/mL catalase, 20 mg/mL glucose oxidase) supplemented with 2 mM ATP and 7 μM unlabeled tubulin (gift of R. Ohi, University of Michigan)] was added to the flow chamber and images of microtubules in 8 fields of view were obtained every 5 min (5,10,15,20 min time points) using a Ti-E/B inverted TIRF microscope (Nikon) equipped with a 100X 1.49 N.A. oil immersion TIRF objective (Nikon), three 20 mW diode lasers (488 nm, 561 nm and 640 nm), and EMCCD detector (iXon X3DU897, Andor). The total length of microtubules in each field of view was measured using Fiji/ImageJ (National Institutes of Health) and the measurements were summed across 8 fields of view for each time point. Data are from three independent experiments.

### Negative staining and electron microscopy

Taxol-stabilized GDP microtubules, GMPCPP microtubules, and GDP-microtubules were prepared as described [76]. For taxol-stabilized GDP microtubules, 10 µl of glycerol-free tubulin (Cytoskeleton) at 100 µM was incubated for 1 hr at 37°C in BRB80 supplemented with 1mM GTP and 10% DMSO. 20 µl of BRB80 with taxol (warmed up to 37°C) was added to bring the final taxol concentration in the mixture to 20µM. The microtubules were incubated at room temperature overnight and then pelleted through100µl cushion (BRB80, 60% glycerol, and 20 μM Taxol) at 50,000 × g for 45 min at 30°C in a TLA100 rotor. The microtubule pallet was washed twice with BRB80, 20µM taxol, and resuspended in 50µl BRB80, 20 μM Taxol. For GMPCPP microtubules, 20 µl of 100 µM glycerol-free tubulin in BRB80 supplemented with 1mM GMPCPP and 1mM DTT was incubated on ice for 5-10 min and then transferred to 37 °C for 1 hr. The microtubules were pelleted by centrifugation at 50,000 × g for 10 min at 30°C, resuspended in 50 μl ice-cold BRB80 + 1mM DTT and incubated on ice for 30 min with pipetting up and down every five min to depolymerize microtubules. GMPCPP at final concentration of 1 mM was added to the depolymerized tubilin mixture and incubated on ice for another 10 min, then incubated overnight at 37 °C. GMPCPP-microtubules were pelleted through 100µl cushion buffer (BRB80, 60% glycerol) at 50,000 × g for 45 min at 30°C, washed twice with BRB80, and resuspended in 50 µl BRB80. For GDP-microtubules, 20 µl of 100 µM glycerol-free tubulin was incubated at 37 °C for 1 hr in BRB80 supplemented with 1mM GTP and 10% DMSO. The microtubules were pelleted by centrifugation through a cushion (BRB80, 60% glycerol, and 1mM GTP) at 50,000 × g for 45 min at 30°C, washed twice in BRB80, 1mM GTP and 10% DMSO, and resuspended in the same buffer.

Copper grids with Formvar carbon film (FCF400-CU, EMS) were cleaned using PELCO easiGlow™ at 5 mAmp for 30 seconds. 3µl of each microtubule at 1µM (based on original tubulin dimer) in its resuspension buffer was incubated on the glow-discharged grid for 1 minute. Then 5 µl of 25nM KIF5C(1-560)-WT or KIF5C(1-560)-Δ6 in BRB80 buffer supplemented with 10 mM ATP was added to the microtubules on the grid and incubated at 25 °C, 100% humidity for another 5 min. Temperature and humidity were set using a water bath or virtobot instrument. The extra solution was blotted from the grids using calcium-free Whatman filter papers and followed by negative staining with 0.75% uranyl formate. Imagining was performed at room temperature with Morgagni transmission electron microscope (FEI) equipped with a CCD camera and an acceleration voltage of 100 kV. Images were collected at 22,000 magnification (2.1 Å/pixel).

### Statistical Analysis

Statistical analyses were performed and Graphs were generated using Prism software (GraphPad 8.0.0). Comparisons were carried out using a two-tailed *t* test.

## Acknowledgements

We are grateful to David Sept, Ryoma Ohi, Morgan DeSantis and members of their laboratories for helpful discussions and reagents. This work was supported by grants from the National Institutes of Health to KJV (R01GM070862, R35GM131744) and a grant from National Science Foundation to MJL (1330792). BB was supported by the Cellular and Molecular Biology Training Grant T32-GM007315 from the National Institutes of Health, a Graduate Research Fellowship (DGE 1256260) from the National Science Foundation, an EDGE Fellowship from the Endowment of Basic Sciences at the University of Michigan Medical School, and a Rackham Predoctoral Fellowship from the Horace H. Rackham School of Graduate Studies at the University of Michigan. DNR was supported by the National Science Foundation Graduate Research Fellowship Program under grant no. 1445197.

## Movies

**Movie 1. Expression of KIF5C(1-560)-Δ6 leads to buckling and breakage of microtubules in cells**. Live-cell imaging of COS-7 cells expressing EGFP α-tubulin (green) and Halo-Flag-tagged KIF5C(1-560)-Δ6 labeled with JF552 (magenta). A microtubule buckles at the cell periphery and then breaks within the buckled region (scale bar, 5 μm). Images were acquired every 200 ms and playback rate is 80 fps.

**Movie 2. Expression of KIF5C(1-560)-Δ6 leads to knotting and breakage of microtubules in cells**. Live-cell imaging of COS-7 cells expressing EGFP α-tubulin (green) and Halo-Flag-tagged KIF5C(1-560)-Δ6 labeled with JF552 (magenta). A microtubule extends from a region of knotted microtubules and then breaks near the knot while its plus end remains paused at the cell periphery (scale bar, 5 μm). Images were acquired every 200 ms and playback rate is 80 fps.

**Movie 3. Expression of KIF5C(1-560)-Δ6 leads to bending, looping, buckling, and breakage of microtubules in cells**. Live-cell imaging of COS-7 cells expressing EGFP α-tubulin (green) and Halo-Flag-tagged KIF5C(1-560)-Δ6 labeled with JF552 (magenta). Images were acquired every 5 s and playback rate is 2 fps. Scale bar, 5 μm.

---

**Figure S1.**
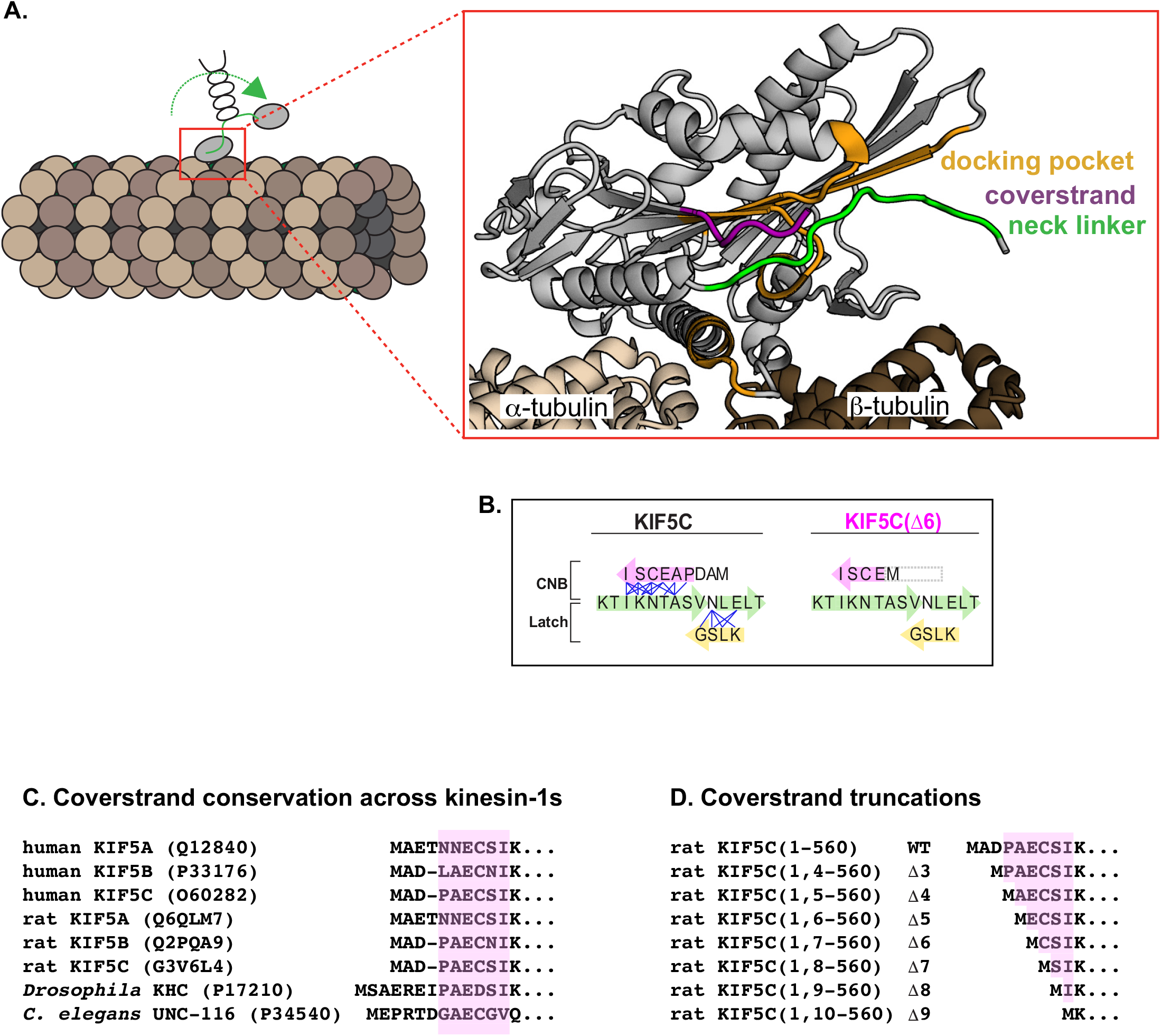
Neck linker docking of kinesin-1. (A) Schematic of key structural elements involved in neck linker (NL) docking. The kinesin-1 motor domain (RnKIF5C) in the ATP-bound, post-power stroke state is shown as a cartoon representation (PDB 4HNA) with secondary structure elements: coverstrand (β0, purple), neck linker (β9 and β10, green), and docking pocket formed by β7 and the loop between β1 and β3 (yellow). (B) Schematic of NL docking. The first half of the NL (β9, green) interacts with the coverstrand (β0, magenta) to form the cover-neck bundle (CNB). The second half of the NL (β10, green) interacts with β7 of the docking pocket (yellow) for NL latching. Residue-residue contacts for NL docking are depicted as blue lines (Budaitis et al., 2019). The coverstrand residues missing in the Δ6 truncation are indicated by a grey box. (C) Amino acid sequence alignment showing the coverstrand conservation across members of the kinesin-1 family. (D) Schematic of truncations initiated to probe the length-dependence of the coverstrand for force generation.

**Figure S2.**
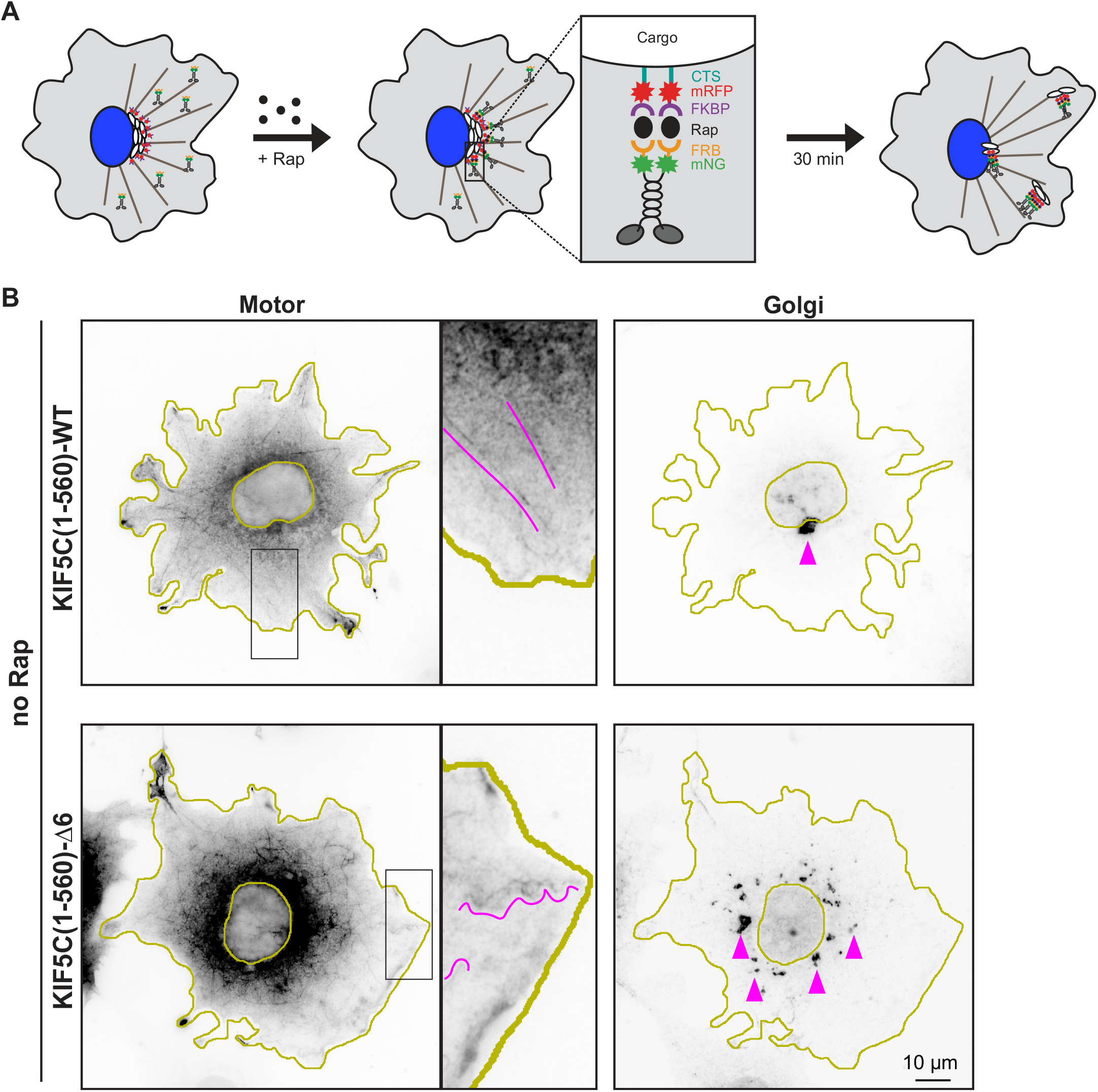
Expression of KIF5C(1-560)-Δ6 results in dispersion of the Golgi complex. (A) Schematic of the inducible motor recruitment assay. A kinesin motor tagged with monomeric NeonGreen (mNG) and an FRB domain (KIF5C-mNG-FRB) is co-expressed with a cargo targeting sequence (CTS) tagged with monomeric red fluorescent protein (mRFP) and FKBP domain (CTS-mRFP-FKBP) in COS-7 cells. Addition of rapamycin (+Rap) causes heterodimerization of the FRB and FKBP domains and recruitment of motors to the cargo membrane. Recruitment of active motors drives cargo dispersion to the cell periphery. (B) Representative images of motor and Golgi localization in COS-7 cells co-transfected with a plasmid encoding for the expression of a Golgi-targeted GMAP210p-RFP-FKBP and (top) KIF5C(1-560)-WT-mNG-FRB or (bottom) KIF5C(1-560)-Δ6-mNG-FRB motors. The cells were fixed and stained with an antibody against the Golgi protein giantin. In cells expressing KIF5C(1-560)-WT, the Golgi localizes in a characteristic tight complex adjacent to the nucleus. In contrast, in cells expressing KIF5C(1-560)-Δ6, the Golgi complex is dispersed within the cytoplasm. Magenta lines denote motor decoration of (top) straight or (bottom) highly curved structures in the cell; magenta arrowheads indicate Golgi elements; yellow lines indicate the nucleus and cell periphery. Scale bar, 10 μm.

**Figure S3.**
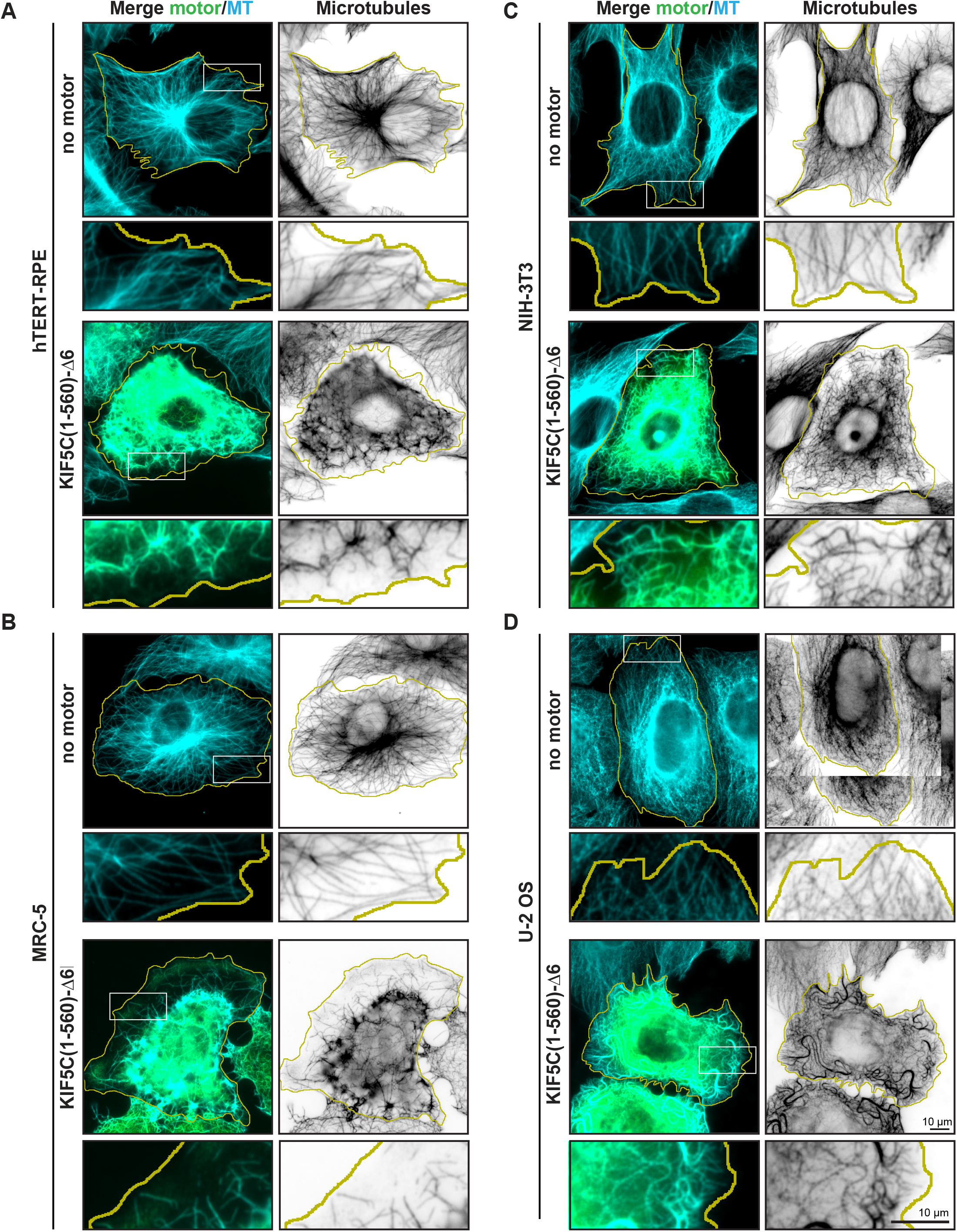
Expression of KIF5C(1-560)-Δ6 in multiple cell lines. (A-D) Representative images of the microtubule network in (A) human hTERT-RPE, (B) human MRC-5, (C) mouse NIH-3T3, and (D) human U-2 OS cells expressing 3xmCit-tagged KIF5C(1-560)-Δ6. Scale bars, 10 μm.

**Figure S4.**
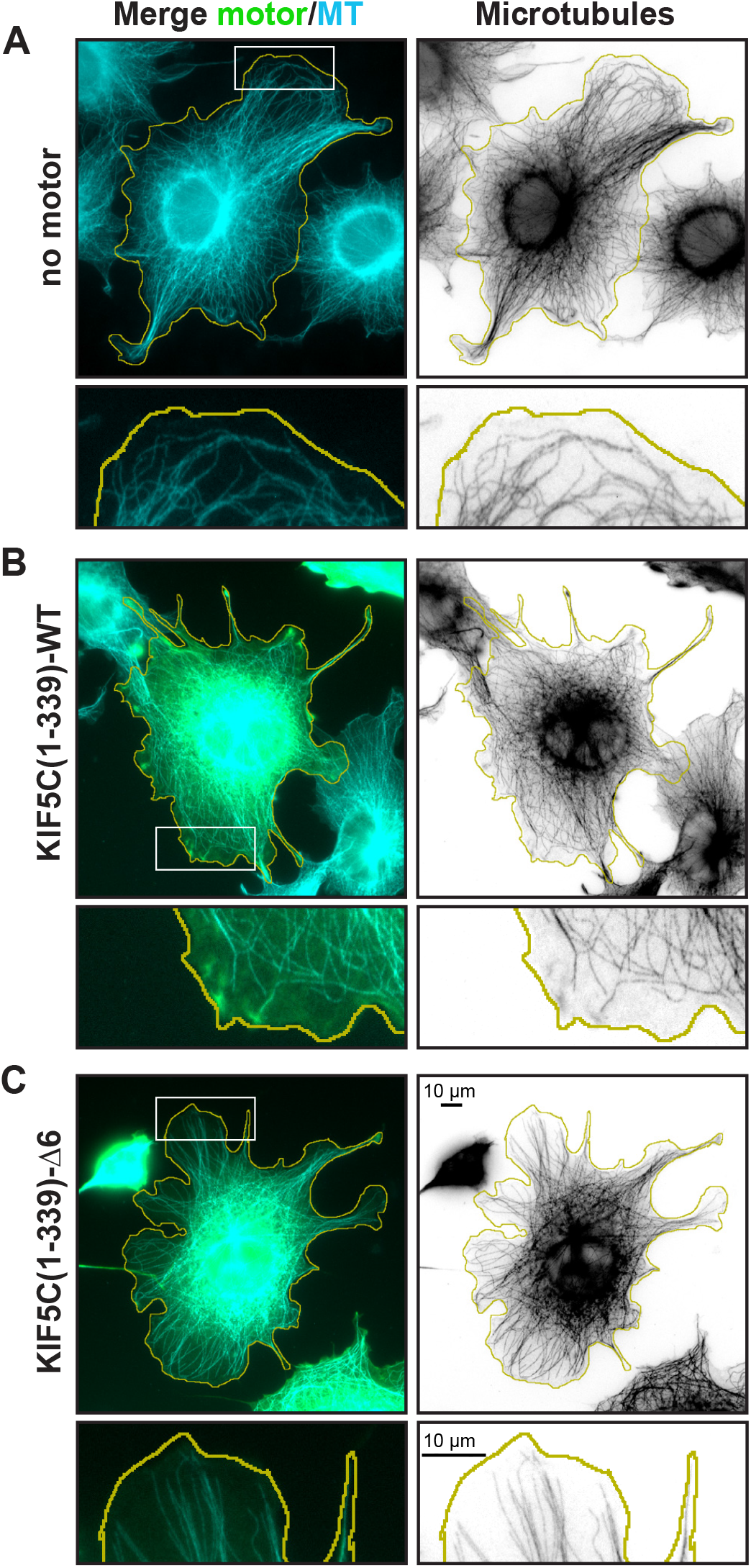
Expression of monomeric KIF5C(1-339)-Δ6 does not result in microtubule breakage and fragmentation. (A-C) Representative images of the microtubule network in (A) untransfected COS-7 cells or COS-7 cells transfected with plasmids encoding for the expression of 3xmCit-tagged (B) monomeric KIF5C(1-339)-WT or (C) monomeric KIF5C(1-339)-Δ6 mutant. Yellow lines indicate the periphery of each cell; white boxes indicate regions presented at higher magnification below each image. Scale bars, 10 μm.

**Figure S5.**
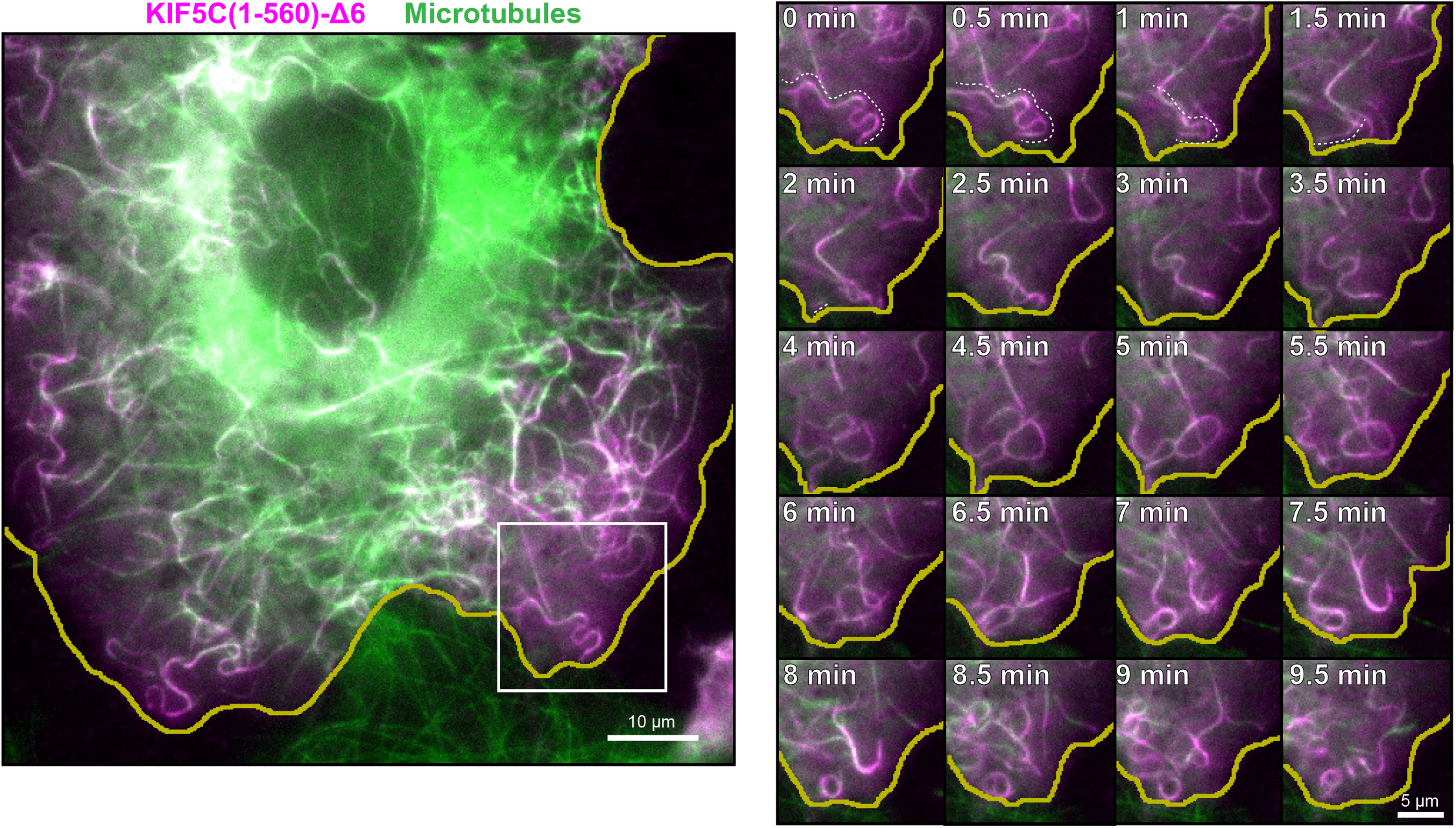
Expression of KIF5C(1-560)-Δ6 results in microtubule bending, looping, buckling and breakage. Live-cell imaging of COS-7 cells expressing EGFP-α-tubulin (green) and Halo-Flag-tagged KIF5C(1-560)-Δ6 labeled with JF552 (magenta). (Left) still image and (right) frames from white boxed region. White dashed line highlights a microtubule that buckles and breaks followed by depolymerization of the fragment from the minus end. Yellow lines indicate the periphery of the cell. Scale bars, 10 μm (left) and 5 μm (right).

**Figure S6.**
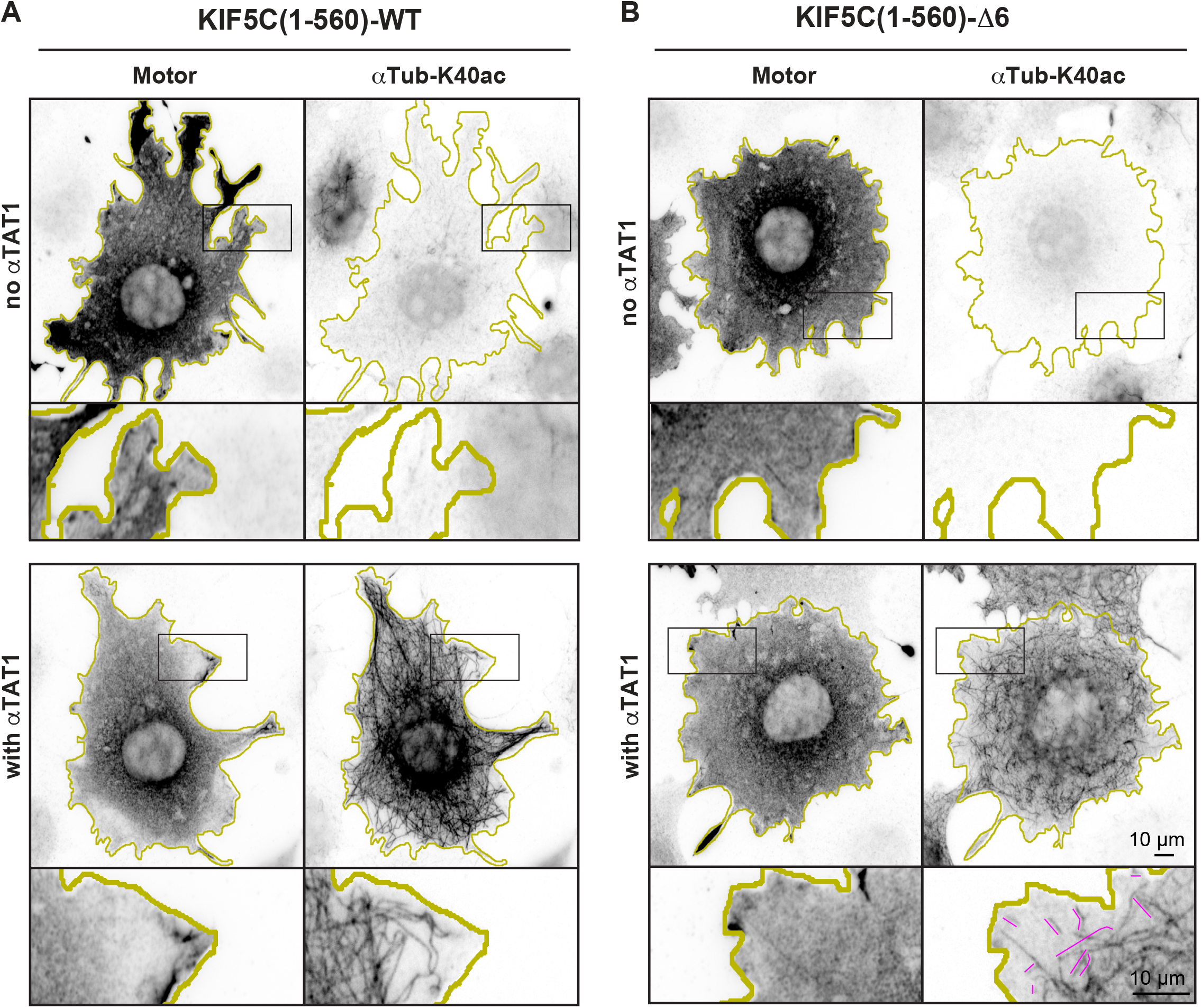
Expression of αTAT1 drives increased microtubule acetylation. (A,B) Representative images of the microtubule acetylation in COS-7 cells transfected with plasmids encoding mCit-αTAT1 and either (A) KIF5C(1-560)-WT or (B) KIF5C(1-560)-Δ6. The cells were fixed and stained with an antibody against αTub-K40ac. Yellow lines indicate the periphery of each cell; black boxes indicate regions shown at higher magnification below each image; magenta lines indicate fragmented microtubules. Scale bars, 10 μm.

**Figure S7.**
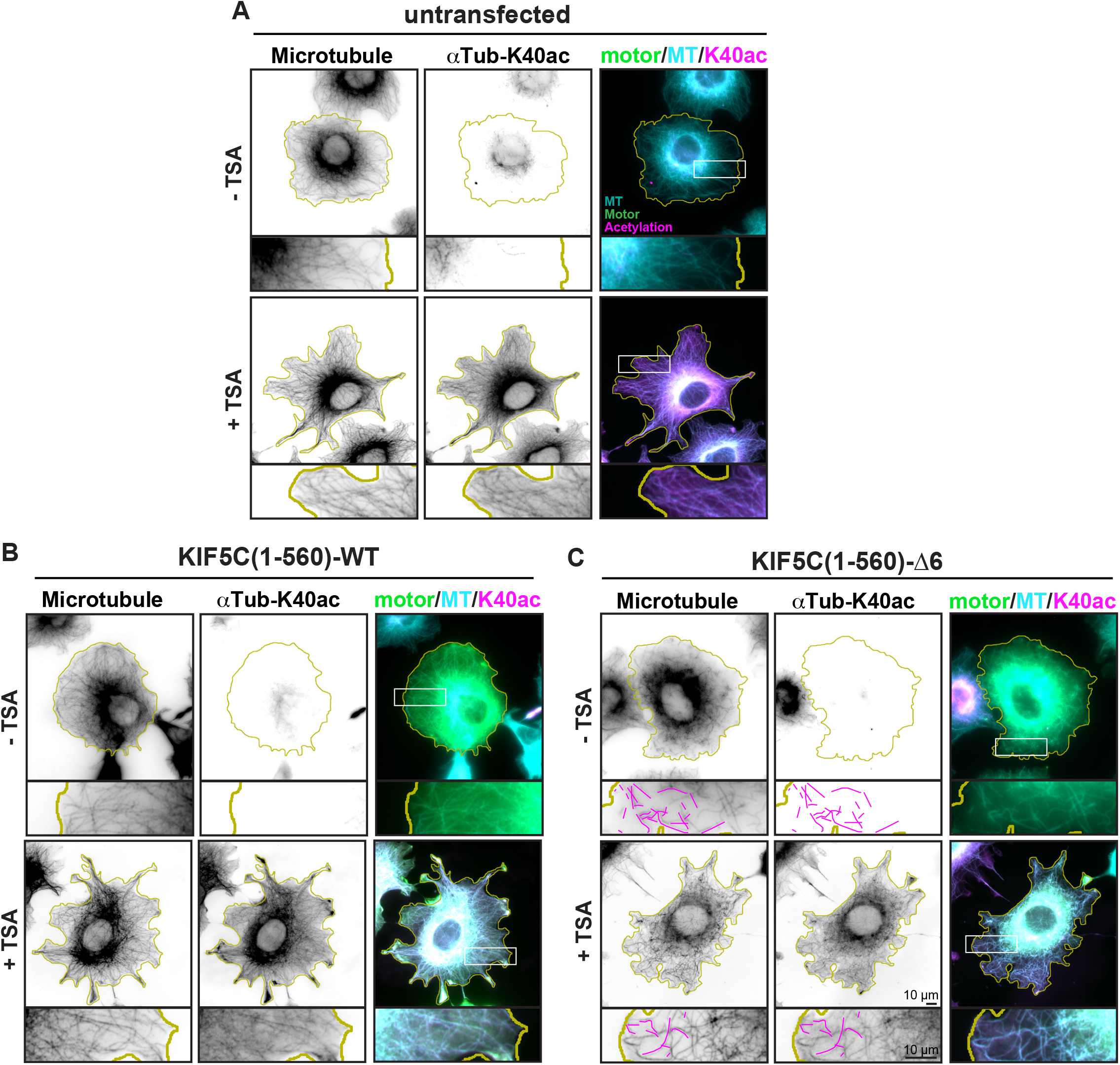
TSA treatment results in increased microtubule acetylation but does not protect the microtubule network from KIF5C(1-560)-Δ6 destruction. (A-C) Representative images. COS-7 cells were (A) untransfected or transfected with plasmids encoding for the expression of (B) KIF5C(1-560)-WT or (C) KIF5C(1-560)-Δ6. 16 hr later, the cells were treated with the deacetylase inhibitor trichostatin A (TSA) for 4 hr and then the cells were fixed and stained with antibodies against β-tubulin and αTub-K40ac. Yellow lines indicate the periphery of each cell; white boxes indicate regions shown at higher magnification below each image; magenta lines indicate fragmented microtubules. Scale bars, 10 μm.

**Figure S8.**
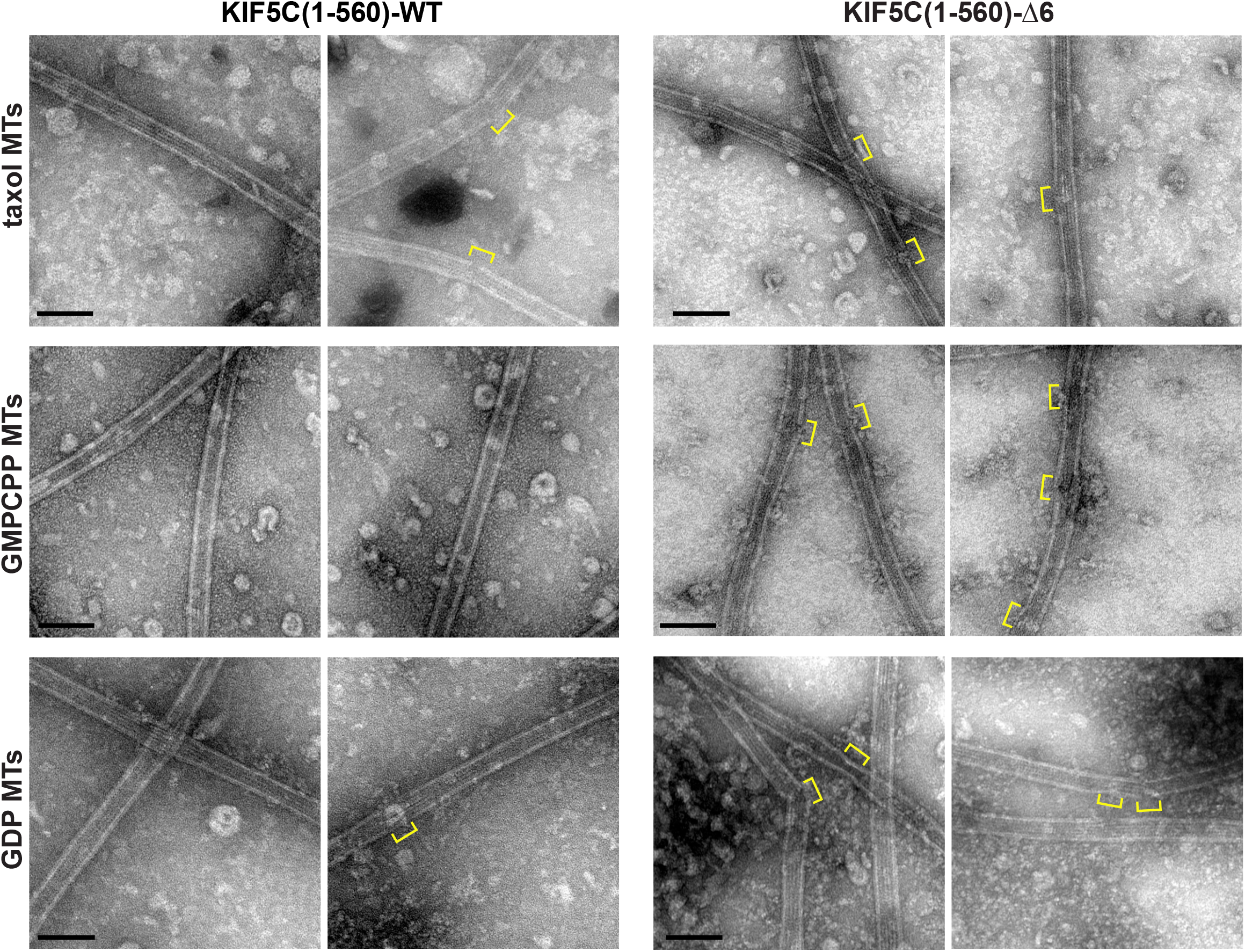
Additional images related to Figure 7. Taxol-stabilized microtubules (taxol MTs), GMPCPP-tubulin microtubules (GMPCPP MTs), or GDP-tubulin microtubules (GDP MTs) were incubated on grids with purified Halo-Flag-tagged KIF5C(1-560)-WT or KIF5C(1-560)-Δ6 motors for 5 min before blotting and staining with uranyl formate. Yellow brackets, sites of lattice damage. Scale bars,100 nm.

